# A Guide to Robust Statistical Methods in Neuroscience

**DOI:** 10.1101/151811

**Authors:** Rand R. Wilcox, Guillaume A. Rousselet

## Abstract

There is a vast array of new and improved methods for comparing groups and studying associations that offer the potential for substantially increasing power, providing improved control over the probability of a Type I error, and yielding a deeper and more nuanced understanding of neuroscience data. These new techniques effectively deal with four insights into when and why conventional methods can be unsatisfactory. But for the non-statistician, the vast array of new and improved techniques for comparing groups and studying associations can seem daunting, simply because there are so many new methods that are now available. The paper briefly reviews when and why conventional methods can have relatively low power and yield misleading results. The main goal is to suggest some general guidelines regarding when, how and why certain modern techniques might be used.

## 1 Introduction

The typical introductory statistics course covers classic methods for comparing groups (e.g., Students t-test, the ANOVA F test and the Wilcoxon–Mann–Whitney test) and studying associations (e.g., Pearsons correlation and least squares regression). The two-sample Students t-test and the ANOVA F test assume that sampling is from normal distributions and that the population variances are identical, which is generally known as the homoscedasticity assumption. When testing hypotheses based on the least squares regression estimator or Pearsons correlation, similar assumptions are made. (Section 2.2 elaborates on the details.) An issue of fundamental importance is whether violating these assumptions can have a serious detrimental impact on two key properties of a statistical test: the probability of a false positive, also known as a Type I error, and power, the probability of detecting true differences among groups and a true association among two or more variables. There is the related issue of whether conventional methods provide enough detail regarding how groups differ as well as the nature of true association.

There are a variety of relatively well-known techniques for dealing with non-normality and unequal variances. For example, use a rank based method. However, by modern standards, these methods are relatively ineffective for reasons reviewed in section 3. More effective techniques are indicated in section 4.

The good news is that when comparing groups that have non-normal but identical distributions, control over the Type I error probability is, in general, reasonably good when using conventional techniques. But if the groups differ, there is now a vast literature indicating that under general conditions, power can be relatively poor. In practical terms, important differences among groups might be missed (e.g., Wilcox, 2017a, b, c). Even when the normality assumption is true, but the population variances differ (called heteroscedasticity), power can be adversely impacted when using the ANOVA F.

Similar concerns arise when dealing with regression. Conventional methods, including rank-based techniques, perform well, in terms of controlling the probability of a Type I error, when there is no association. But when there is an association, conventional methods, including rank-based techniques (e.g., Spearmans rho and Kendalls tau) can have a relatively low probability of detecting an association relative to modern methods developed during the last thirty years.

Practical concerns regarding conventional methods stem from four major insights (e.g., Wilcox, 2017a, c). These insights can be briefly summarized as follows.

- The central limit and skewed distributions: much larger sample sizes might be needed to assume normality than is generally recognized.
- There is now a deeper understanding of the role of outliers and how to deal with them. Some seemingly obvious strategies for dealing with outliers, based on standard training, are known to be highly unsatisfactory for reasons outlined later in the paper.
- There is a substantial literature indicating that methods that assume homoscedasticity (equal variances) can yield inaccurate results when in fact there is heteroscedasticity, even when the sample sizes are quite large.
- When dealing with regression, curvature refers to situations where the regression line is not straight. There is now considerable evidence that curvature is a much more serious concern than is generally recognized.

Robust methods are typically thought of as methods that provide good control over the probability of a Type I error. But today they deal with much broader issues. In particular, they are designed to deal with the problems associated with skewed distributions, outliers, heteroscedasticity and curvature that were outlined above.

One of the more fundamental goals among robust methods is to develop techniques that are not overly sensitive to very small changes in a distribution. For instance, a slight departure from normality should not destroy power. This rules out any method based on the mean and variance (e.g., Staudte & Sheather, 1990; Wilcox, 2017a, b). Section 2.3 illustrates this point.

Many modern robust methods are designed to have nearly the same amount of power as conventional methods under normality, but they continue to have relatively high power under slight departures from normality where conventional techniques based on means perform poorly. There are other fundamental goals, some of which are relevant regardless of how large the sample sizes might be. But an effective description of these goals goes beyond the scope of this paper. For present purposes, the focus is on achieving relatively high power.

Another point that should be stressed has to do with standard power analyses. A common goal is to justify some choice for the sample sizes prior to obtaining any data. Note that in effect, the goal is to address a statistical issue without any data. Typically this is done by assuming normality and homoscedasticity, which in turn can suggest that relatively small sample sizes provide adequate power when using means. A practical concern is that violating either of these two assumptions can have a tremendous impact on power when attention is focused exclusively on comparing means. Section 2.1 illustrates this concern when dealing with measures of central tendency. Similar concerns arise when dealing with least squares regression and Pearson’s correlation. These concerns can be mitigated by using recently developed robust methods summarized here as well as in Wilcox (2017a, c).

There is now a vast array of new and improved methods that effectively deal with known concerns associated with classic techniques (e.g., Maronna et al., 2006; Heritier et al., 2009; Wilcox, 2017a, b, c). They include substantially improved methods for dealing with all four of the major insights previously listed. Perhaps more importantly, they can provide a deeper, more accurate and more nuanced understanding of data as will be illustrated in section 5.

For books focused on the mathematical foundation of modern robust methods, see Hampel et al. (1986), Huber and Ronchetti (2009), Maronna et al. (2006), and Staudte and Sheather (1990). For books focused on applying robust methods, see Heritier et al. (2009) and Wilcox (2017a, c). From an applied point of view, the difficulty is not finding a method that effectively deals with violations of standard assumptions. Rather, for the non-statistician, there is the difficulty of navigating through the many alternative techniques that might be used. This paper is an attempt to deal with this issue by providing a general guide regarding when and how modern robust methods might be used when comparing two or more groups. When dealing with regression, all of the concerns associated with conventional methods for comparing groups remain and new concerns are introduced. A few issues related to regression and correlations are covered here, but it is stressed that there are many other modern advances that have practical value. Readers interested in regression are referred to Wilcox (2017a, c).

A few general points should be stressed. First, if robust methods, such as modern methods based on the median described later in this paper, give very similar results to conventional methods based on means, this is reassuring that conventional methods based on the mean are performing relatively well in terms of Type I errors and power. But when they differ, there is doubt about the validity of conventional techniques. In a given situation, conventional methods might perform well in terms of controlling the Type I error probability and providing reasonably high power. But the best that can be said is that there are general conditions where conventional methods do indeed yield inaccurate inferences. A particular concern is that they can suffer from relatively low power in situations where more modern methods have relatively high power. More details are provided in sections 3 and 4.

Second, the choice of method can make a substantial difference in our understanding of data. One reason is that modern methods provide alternative and interesting perspectives that more conventional methods do not address. A complication is that there is no single method that dominates in terms of power or providing a deep understanding of how groups compare. The same is true when dealing with regression and measures of association. The reality is that several methods might be needed to address even what appears as a simple problem, for instance comparing two groups.

There is, of course, the issue of controlling the probability of one or more Type I errors when multiple tests are performed. There are many modern improvements for dealing with this issue (e.g., Wilcox, 2017a, c). And another strategy is to put more emphasis on exploratory studies. One could then deal with the risk of false positive results by conducting a confirmatory study aimed at determining whether significant results in an exploratory study can be replicated (Wagenmakers et al., 2012). Otherwise, there is the danger of missing important details regarding how groups compare. One of the main messages here is that despite the lack of a single method that dominates, certain guidelines can be offered regarding how to analyze data.

Modern methods for plotting data can be invaluable as well (e.g., Rousselet, et al., 2017; Rousselet et al., 2016; Weissgerber et al., 2016) In particular, they can provide important perspectives beyond the common strategy of using error bars. Complete details go beyond the scope of this paper, but section 5 illustrates some of the more effective plots that might be used.

The paper is organized as follows. Section 2 briefly reviews when and why conventional methods can be highly unsatisfactory. This is necessary in order to appreciate modern technology and because standard training typically ignores these issues. Efforts to modernize basic training have been made (e.g., Field et al., 2012; Wilcox, 2017b, c). And a 2016 special issue of the *American Statistician* (volume 69, number 4) was aimed at encouraging instructors to modernize their courses. (This special issue touches on a broader range of topics than those discussed here.) Some neuroscientists are trained in a manner that takes into account modern insights relevant to basic principles. But it is evident that most are not. Section 3 reviews the seemingly more obvious strategies aimed at salvaging standard techniques, the point being that by modern standards they are relatively ineffective and cannot be recommended. Moreover, certain strategies are not technically sound. Section 3 also provides an indication of how concerns regarding conventional methods are addressed using more modern techniques. Section 4 describes strategies for comparing two independent or dependent groups that take modern advances into account. Included are some methods aimed at comparing correlations as well as methods designed to determine which independent variables are most important. Section 5 illustrates modern methods using data from several studies.

## 2 Insights Regarding Conventional Methods

This section elaborates on the concerns with conventional methods for comparing groups and studying associations stemming from the four insights previously indicated.

### 2.1 Skewed Distributions

A skewed distribution simply refers to a distribution that is not symmetric about some central value. An example is shown in Figure 1**A**. Such distributions occur naturally. An example relevant to the neurosciences is given in section 5.2. Skewed distributions are a much more serious problem for statistical inferences than once thought due to insights regarding the central limit theorem.

**Figure 1:**
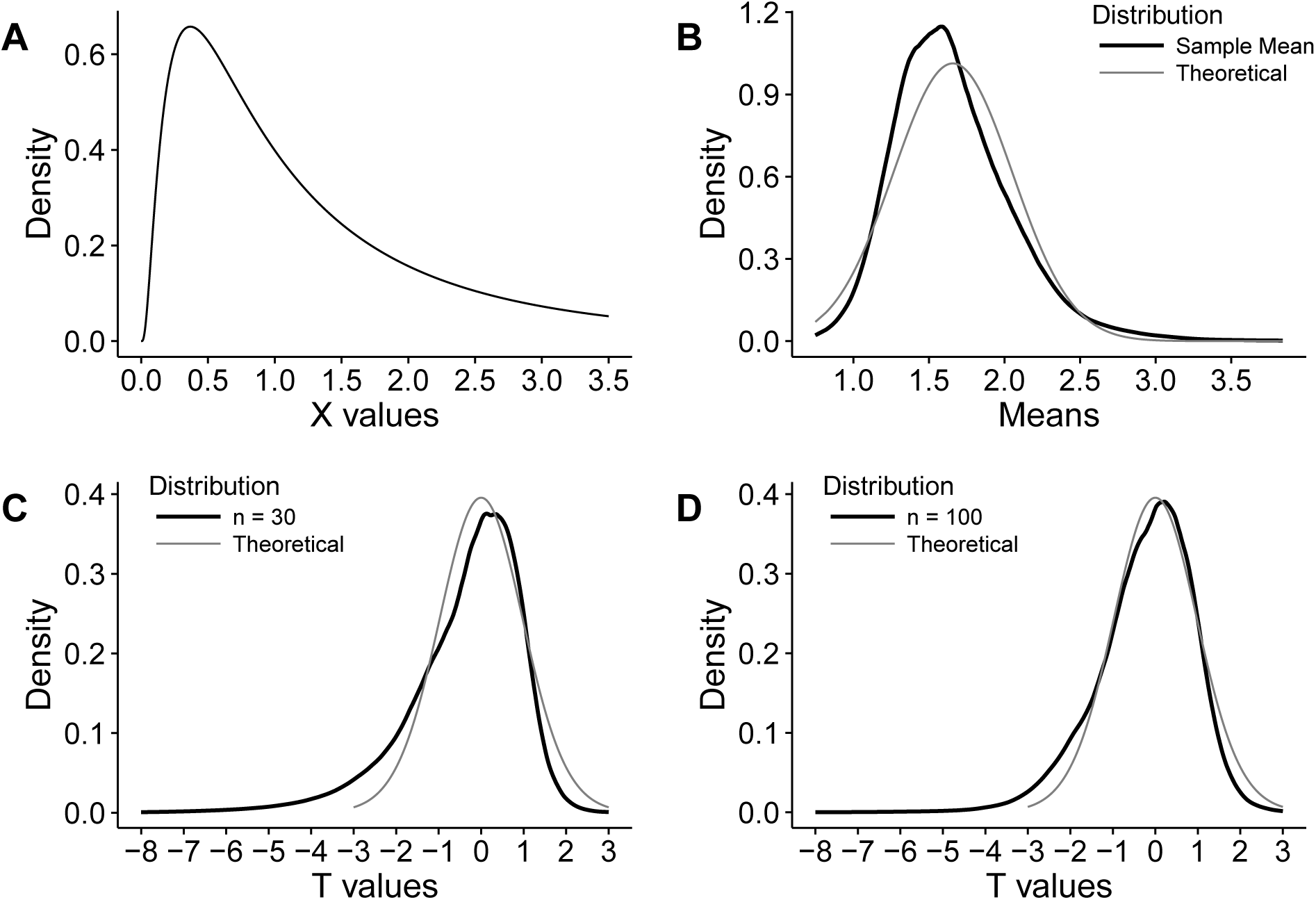
Panel **A** illustrates an example of skewed distribution. Panel **B** illustrates the distribution of the sample mean under normality (the dashed line), *n* = 30, and the actual distribution based on a simulation. Each sample mean was computed based on 30 observations randomly sampled from the distribution shown in **A.** Panels **C** and **D** compare the theoretical *T* distribution with 29 degrees of freedom to distributions of 5000 *T* values. Again, the *T* values were computed from observations sampled from the distribution in **A.**

Consider the one-sample case. Conventional wisdom is that with a relatively small sample size, normality can be assumed under random sampling. An implicit assumption was that if the sample mean has, approximately, a normal distribution, then Students t-test will perform reasonably well. It is now known that this is not necessarily the case as illustrated, for example, in Wilcox (2017a, b).

This point is illustrated here using a simulation that is performed in the following manner. Imagine that data are randomly sampled from the distribution shown in Figure 1**A** (a lognormal distribution) and the mean is computed based on a sample size of *n* = 30. Repeating this process 5000 times, the thick black line in Figure 1**B** shows a plot of the resulting sample means; the thin gray line is the plot of the means when sampling from a normal distribution instead. The distribution of *T* values for samples of *n* = 30 is indicated by the thick black line in Figure 1**C**; the thin gray line is the distribution of *T* values when sampling from a normal distribution. As can be seen, the actual *T* distribution extends out much further to the left compared to the distribution of *T* under normality. That is, in the current example, sampling from a skewed distribution leads to much more extreme values than expected by chance under normality, which in turn results in more false positive results than expected when the null hypothesis is true.

Suppose the goal is to test some hypothesis at the 0.05 level. Bradley (1978) suggests that as a general guide, control over the probability of a Type I error is minimally satisfactory if the actual level is between 0.025 and 0.075. When we test at the 0.05 level, we expect 5% of the t-tests to be significant. However, when sampling from the skewed distribution considered here, this is not the case: the actual Type I error probability is approximately 0.111.

Figure 1**D** shows the distribution of *T* when *n* = 100. Now the Type I error probability is approximately 0.082, again when testing at the 0.05 level. Based on Bradleys criterion, a sample size of about 130 or larger is required. Bradely (1978) goes on to suggest that ideally, the actual Type I error probability should be between 0.045 and 0.055. Now *n* = 600 is unsatisfactory; the actual level is approximately 0.057. With *n* = 700 the level is approximately 0.055.

Before continuing, it is noted that the median belongs to the class of trimmed means, which refers to the strategy of trimming a specified proportion of the smallest and largest values and averaging the values that remain. For example, if n=10, 10% trimming means that the lowest and highest values are removed and the remaining data are averaged. Similarly, 20% trimming would remove the two smallest and two largest values. Based on conventional training, trimming might seem counterintuitive, but in some situations it can substantially increase our ability to control the Type I error probability, as illustrated next, and trimming can substantially increase power as well for reasons to be explained.

First focus on controlling the probability of a Type I error. (Section 2.3 illustrates one of the reasons methods based on means can have relatively low power.) Figure 2 illustrates the Type I error probability as a function of the sample size, when when using the mean, median and when sampling from the asymmetric (lognormal) distribution in Figure 1**A**. Inferences based on the 20% trimmed mean were made via the method derived by Tukey and McLaughlin (1963). Inferences based on the median were made via the method derived by Hettmansperger and Sheather (2011). (The software used to apply these latter two methods is contained in the R package described at the beginning of section 4.) Also shown is the Type I error probability when sampling from a normal distribution. The gray area indicates Bradley’s criterion.

**Figure 2:**
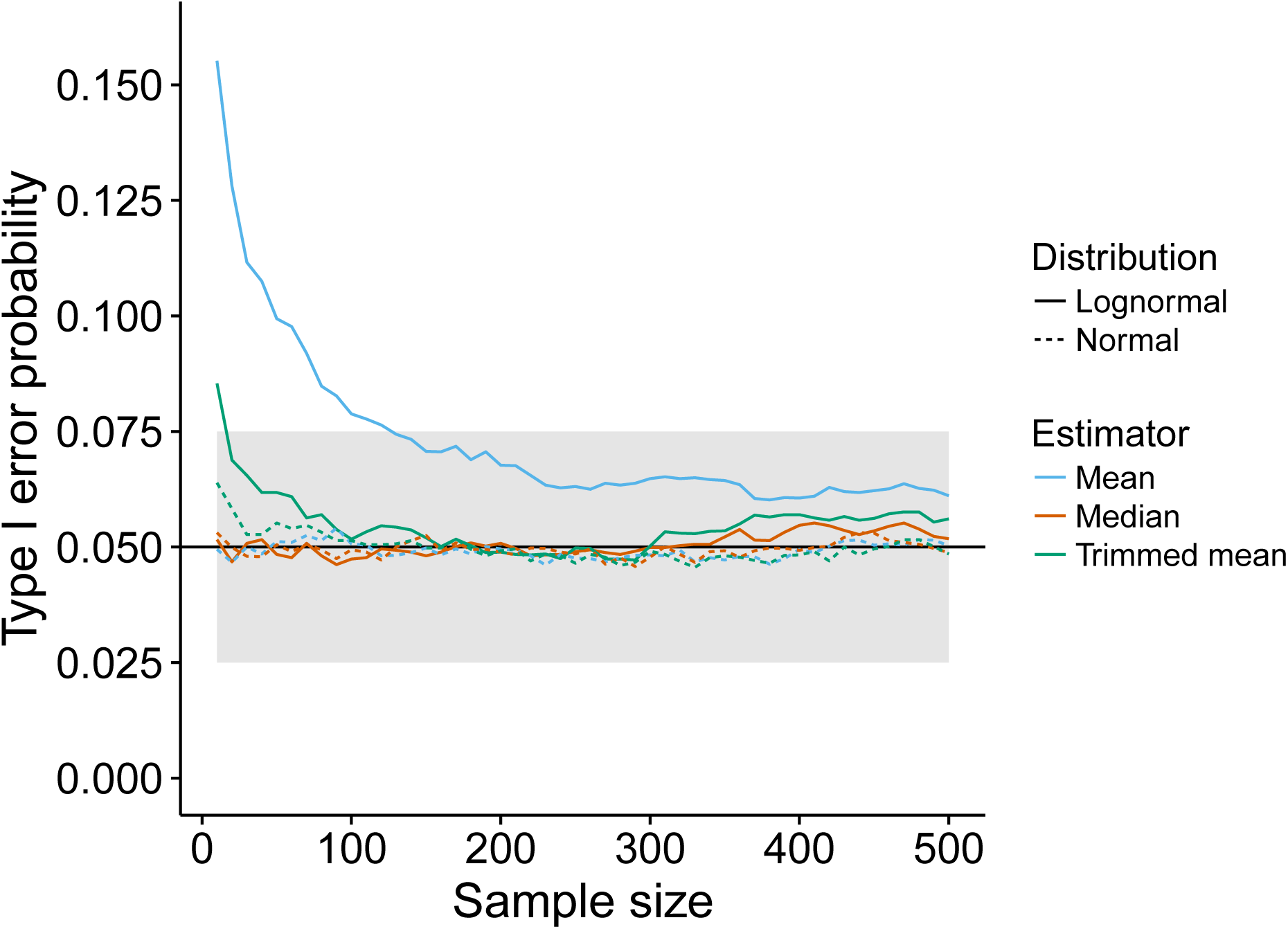
Type I error probability as a function of sample size. The type I error probability was computed by running a simulation with 10,000 iterations. In each iteration, sample sizes from 10 to 500, in steps of 10, were drawn from a normal distribution and a lognormal distribution. For each combination of sample size and distribution, we applied a t-test on the mean, a test of the median, and a t-test on the 20% trimmed mean, all with alpha = 0.05. Depending on the test applied, the mean, median or 20% trimmed mean of the population sampled from was zero. The black horizontal line marks the expected 0.05 type I error probability. The gray area marks Bradley’s satisfactory range. When sampling from a normal distribution, all methods are close to the nominal 0.05 level, except the trimmed mean for very small sample sizes. When sampling is from a lognormal distribution, the mean and the trimmed mean give rise to too many false alarms for small sample sizes. The mean continues to give higher false positive rates than the other techniques even with n=500.

Figure 3 illustrates the association between power and the sample size for the distributions used in Figure 2. As can be seen, under normality, the sample mean is best, followed closely by the 20% trimmed mean. The median is least satisfactory when dealing with a normal distribution, as expected. However, for the asymmetric (lognormal) distribution, the median performs best and the mean performs very poorly.

**Figure 3:**
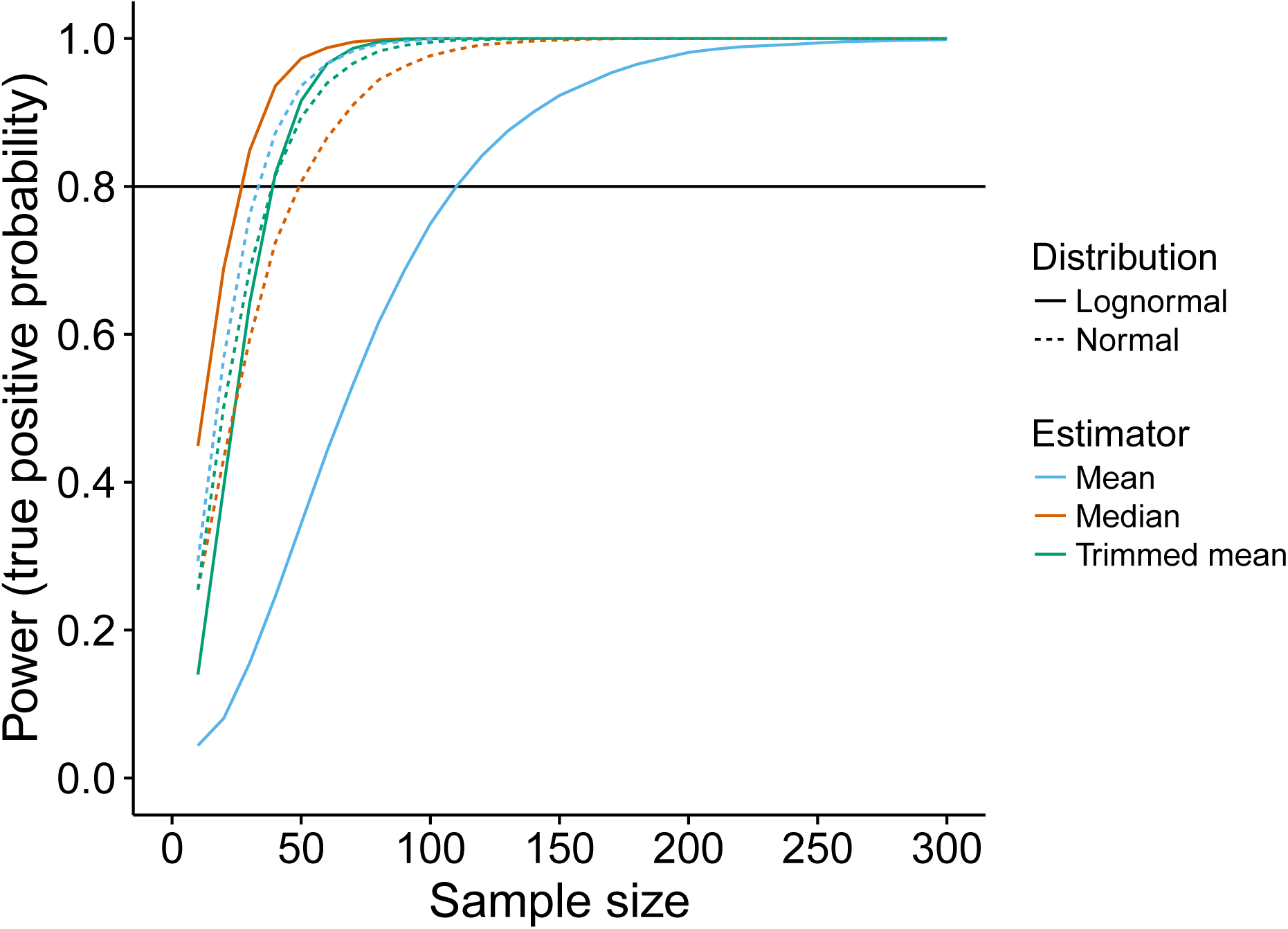
Power as a function of sample size. The probability of a true positive was computed by running a simulation with 10,000 iterations. In each iteration, sample sizes from 10 to 500, in steps of 10, were drawn from a normal distribution and the (lognormal) distribution shown in Figure 1A. For each combination of sample size and distribution, we applied a t-test on the mean, a test of the median, and a t-test on the 20% trimmed mean, all with alpha = 0.05. Depending on the test applied, the mean, median or 20% trimmed mean of the population sampled from was 0.5. The black horizontal line marks the conventional 80% power threshold. When sampling from a normal distribution, all methods require less than 50 observations to achieve 80% power, and the mean appears to have higher power at lower sample size than the other methods. When sampling from a lognormal distribution, power drops dramatically for the mean but not for the median and the trimmed mean. Power for the median actually improves. The exact pattern of results depends on the effect size and the asymmetry of the distribution we sample from, so we strongly encourage readers to perform their own detailed power analyses.

A feature of random samples taken from the distribution in Figure 1**A** is that the expected proportion of points declared an outlier is relatively small. For skewed distributions, as we move toward situations where outliers are more common, a sample size greater than 300 can be required to achieve reasonably good control over the Type I error probability. That is, control over the Type I error probability is a function of both the degree a distribution is skewed and the likelihood of encountering outliers. However, there are methods that perform reasonably well with small sample sizes as indicated in section 4.1.

For symmetric distributions, where outliers tend to occur, the reverse can happen; the actual Type I error probability can be substantially less than the nominal level. This happens because outliers inflate the standard deviation, which in turn lowers the value of *T*, which in turn can negatively impact power. Section 2.3 elaborates on this issue.

In an important sense, outliers have a larger impact on the sample variance than the sample mean, which impacts the t-test. To illustrate this point, imagine the goal is to test *H*_0_: *μ* = 1 based on the following values: 1, 1.5, 1.6, 1.8, 2, 2.2, 2.4, 2.7. Then *T* = 4.69, the p value is *p* = 0.002, and the 0.95 confidence interval = [1.45, 2.35]. Now, including the value 8, the mean increases from 1.9 to 2.58, suggesting at some level there is stronger evidence for rejecting the null hypothesis. However, this outlier increases the standard deviation from 0.54 to 2.1, and now *T* = 2.26 and *p* = 0.054. The 0.95 confidence interval = [–0.033, 3.19].

Now consider the goal of comparing two independent or dependent groups. If the groups have identical distributions, then difference scores have a symmetric distribution and the probability of a Type I error is, in general, less than the nominal level when using conventional methods based on means. Now, in addition to outliers, differences in skewness create practical concerns when using Students t-test. Indeed, under general conditions, the two-sample Students t-test for independent groups is not even asymptotically correct, roughly because the standard error of the difference between the sample means is not estimated correctly (e.g., Cressie & Whitford, 1986). Moreover, Students t-test can be biased. This means that the probability of rejecting the null hypothesis of equal means can be higher when the population means are equal, compared to situations where the population means differ. Roughly, this concern arises because the distribution of *T* can be skewed, and in fact the mean of *T* can differ from zero even though the null hypothesis is true. (For a more detailed explanation, see Wilcox, 2017c, section 5.5.) Problems persist when Students t-test is replaced by Welchs (1938) method, which is designed to compare means in a manner that allows unequal variances. Put another way, if the goal is to test the hypothesis that two groups have identical distributions, conventional methods based on means perform well in terms of controlling the Type I error probability. But if the goal is to compare the population means, and if distributions differ, conventional methods can perform poorly.

There are many techniques that perform well when dealing with skewed distributions in terms of controlling the Type I error probability, some of which are based on the usual sample median (Wilcox, 2017a, c). Both theory and simulations indicate that as the amount of trimming increases, the ability to control over the probability of a Type I error increases as well. Moreover, for reasons to be explained, trimming can substantially increase power, a result that is not obvious based on conventional training. The optimal amount of trimming depends on the characteristics of the population distributions, which are unknown. Currently, the best that can be said is that the choice can make a substantial difference. The 20% trimmed has been studied extensively and often it provides a good compromise between the two extremes: no trimming (the mean) and the maximum amount of trimming (the median).

In various situations, particularly important are inferential methods based on what are called bootstrap techniques. Two basic versions are the bootstrap-t and percentile bootstrap. Roughly, rather than assume normality, bootstrap-t methods perform a simulation using the observed data that yields an estimate of an appropriate critical value and a p-value. So values of *T* are generated as done in Figure 1, only data are sampled, with replacement, from the observed data. In essence, bootstrap-t methods generate data-driven *T* distributions expected by chance if there were no effect. The percentile bootstrap proceeds in a similar manner, only when dealing with a trimmed mean, for example, the goal is to determine the distribution of the sample trimmed mean, which can then be used to compute a p-value and a confidence interval. When comparing two independent groups based on the usual sample median, if there are tied (duplicated) values, currently the only method that performs well in simulations, in terms of controlling the Type I error probability, is based on a percentile bootstrap method (cf. Wilcox, 2017c, Table 5.3.). Section 4.1 elaborates on how this method is performed.

If the amount of trimming is close to zero, the bootstrap-t method is preferable to the percentile bootstrap method. But as the amount of trimming increases, at some point a percentile bootstrap method is preferable. This is the case with 20% trimming (e.g., Wilcox, 2017). It seems to be the case with 10% trimming as well, but a definitive study has not been made.

Also, if the goal is to reflect the typical response, it is evident that the median or even a 20% trimmed mean might be more satisfactory. Using quantiles (percentiles) other than the median can be important as well, for reasons summarized in section 4.2.

When comparing independent groups, modern improvements on the Wilcoxon–Mann– Whitney (WMW) test are another possibility, which are aimed at making inferences about the probability that a random observation from the first group is less than a random observation from the second. (More details are provided in section 3.) Additional possibilities are described in Wilcox (2017a), some of which are illustrated in section 4 of this paper.

In some situations, robust methods can have substantially higher power than any method based on means. But it is not being suggested that robust methods always have more power. This is not the case. Rather, the point is that power can depend crucially on the conjunction of which estimator is used (for instance the mean vs. the median), and how a confidence interval is built (for instance a parametric method or the percentile bootstrap). These choices are not trivial and must be taken into account when analyzing data.

### 2.2 Heteroscedasticity

It has been clear for some time that when using classic methods for comparing means, heteroscedasticity (unequal population variances) is a serious concern (e.g., Brown & Forsythe, 1974). Heteroscedasticity can impact both power and the Type I error probability. The basic reason is that, under general conditions, methods that assume homoscedasticity are using an incorrect estimate of the standard error when in fact there is heteroscedasticity. Indeed, there are concerns regardless of how large the sample size might be. Roughly, as we consider more and more complicated designs, heteroscedasticity becomes an increasing concern.

When dealing with regression, homoscedasticity means that the variance of the dependent variable does not depend on the value of the independent variable. When dealing with age and depressive symptoms, for example, homoscedasticity means that the variation in measures of depressive symptoms at age 23 is the same at age 80 or any age in between as illustrated in Figure 4.

**Figure 4:**
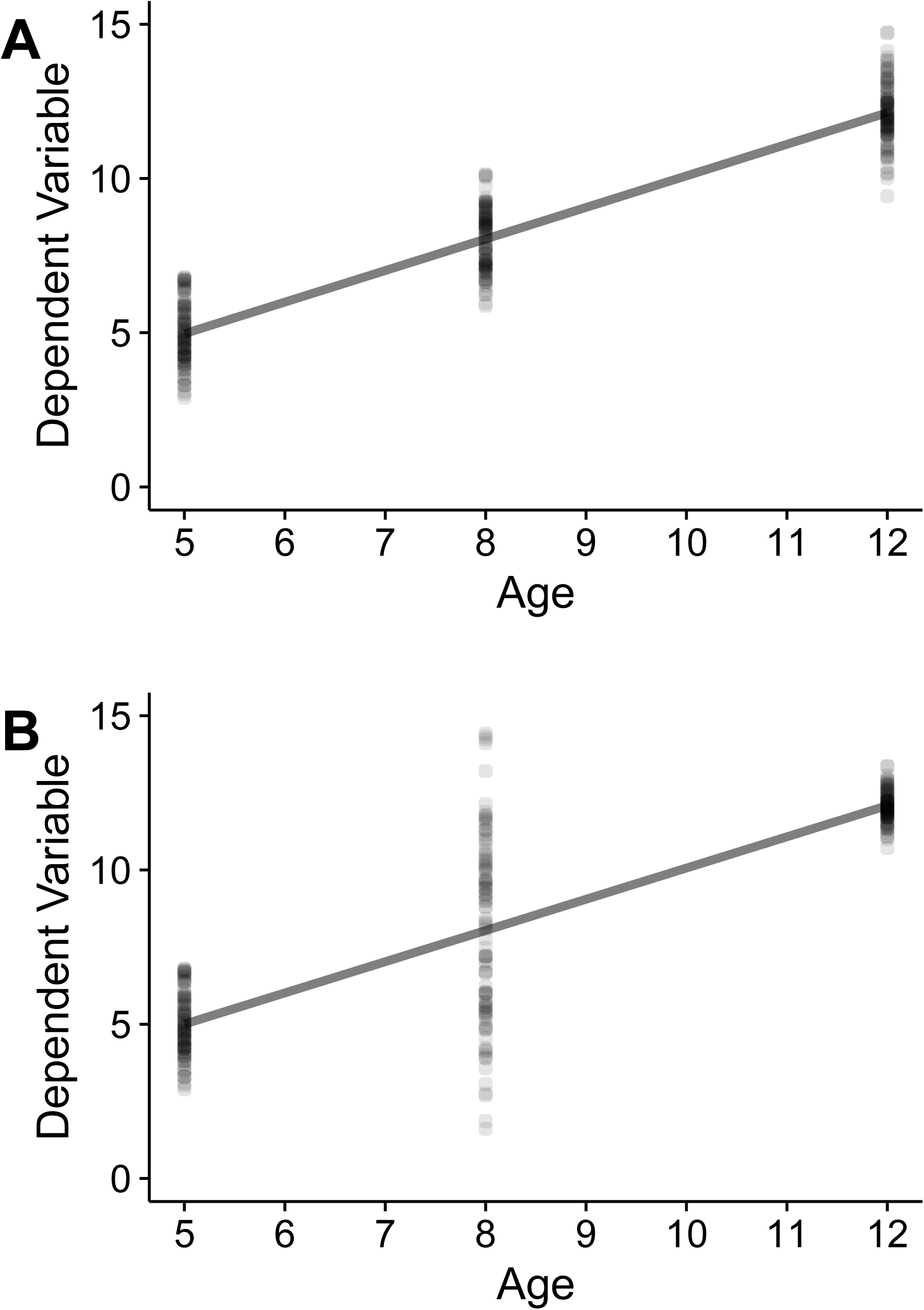
Homoscedasticity and heteroscedasticity. Panel **A** illustrates homoscedasticity. The variance of the dependent variable is the same at any age. Panel **B** illustrates heteroscedasticity. The variance of the dependent variable can depend on the age of the participate.

Independence implies homoscedasticity. So for this particular situation, classic methods associated with least squares regression, Pearson’s correlation, Kendall’s tau and Spearman’s rho are using a correct estimate of the standard error, which helps explain why they perform well in terms of Type I errors when there is no association. That is, when a homoscedastic method rejects, it is reasonable to conclude that there is an association, but in terms of inferring the nature of the association, these methods can perform poorly. Again, a practical concern is that when there is heteroscedasticity, homoscedastic methods use an incorrect estimate of the standard error, which can result in poor power and erroneous conclusions.

A seemingly natural way of salvaging homoscedastic methods is to test the assumption that there is homoscedasticity. But six studies summarized in Wilcox (2017c) found that this strategy is unsatisfactory. Presumably situations are encountered where this is not the case, but it is difficult and unclear how to determine when such situations are encountered.

Methods that are designed to deal with heteroscedasticity have been developed and are easily applied using extant software. These techniques use a correct estimate of the standard error regardless of whether the homoscedasticity assumption is true. A general recommendation is to always use a heteroscedastic method given the goal of comparing measures of central tendency, or making inferences about regression parameters, as well as measures of association.

### 2.3 Outliers

Even small departures from normality can devastate power. The modern illustration of this fact stems from Tukey (1960) and is based on what is generally known as a mixed normal distribution. The mixed normal considered by Tukey means that with probability 0.9 an observation is sampled from a standard normal distribution; otherwise an observation is sampled from a normal distribution having mean zero and standard deviation 10. Figure 5**A** shows a standard normal distribution and the mixed normal discussed by Tukey. Note that in the center of the distributions, the mixed normal is below the normal distribution. But for the two ends of the mixed normal distribution, the tails, the mixed normal lies above the normal distribution. For this reason, the mixed normal is often described as having heavy tails. In general, heavy-tailed distributions roughly refer to distributions where outliers are likely to occur.

**Figure 5:**
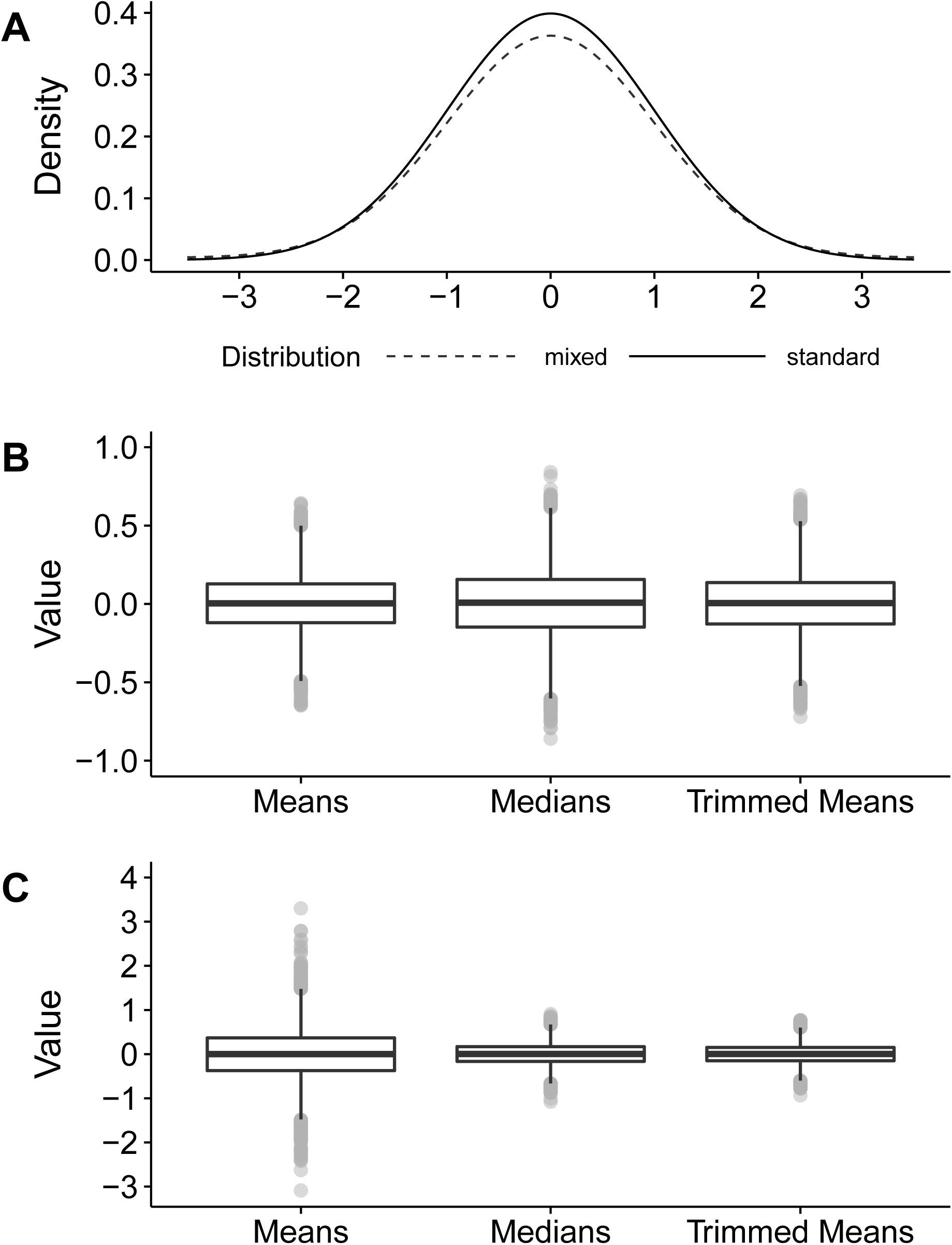
**A.** Density functions for the standard normal distribution (solid line) and the mixed normal distribution (dotted line). These distributions have an obvious similarity, 15 yet the variances are 1 and 10.9. **B.** Boxplots of means, medians and 20% trimmed means when sampling from a normal distribution. **C.** Boxplots of means, medians and 20% trimmed means when sampling from a mixed normal distribution. In panels B and C, each distribution has 10,000 values, and each of these values was obtained by computing the mean, median or trimmed mean of 30 randomly generated observations.

Here is an important point. The standard normal has variance one, but the mixed normal has variance 10.9. That is, the population variance can be overly sensitive to slight changes on the tails of a distribution. Consequently, even slight departure from normality can result in relative poor power when using any method based on the mean.

Put another way, samples from the mixed normal are more likely to result in outliers compared to samples from a standard normal distribution. As previously indicated, outliers inflate the sample variance, which can negatively impact power when using means. Another concern is that they can give a distorted and misleading summary regarding the bulk of the participants.

The first indication that heavy-tailed distributions are a concern stems from a result derived by Laplace about two centuries ago. In modern terminology, he established that as we move from a normal distribution to a distribution more likely to generate outliers, the standard error of the usual sample median can be smaller than the standard error of the mean (Hand, 1998). The first empirical evidence implying that outliers might be more common than what is expected under normality was reported by Bessel (1818).

To add perspective, we computed the mean, median and a 20% trimmed mean based on 30 observations generated from a standard normal distribution. (Again, a 20% trimmed mean removes the 20% lowest and highest values and averages the remaining data.) Then we repeated this process 10,000 times. Boxplots of the results are shown in Figure 5B. Theory tells us that under normality the variation of the sample means is smaller than the variation among the 20% trimmed means and medians, and Figure 1B provides perspective on the extent this is the case.

Now we repeat this process, only data are sampled from the mixed normal in Figure 5**A**. Figure 5**C** reports the results. As is evident, there is substantially less variation among the medians and 20% trimmed means. That is, despite trimming data, the standard errors of the median and 20% trimmed mean are substantially smaller, contrary to what might be expected based on standard training.

Of course, a more important issue is whether the median or 20% trimmed mean ever have substantially smaller standard errors based on the data encountered in research. There are numerous illustrations that this is the case (e.g., Wilcox, 2017a, b, c).

There is the additional complication that the amount of trimming can substantially impact power, and the ideal amount of trimming, in terms of maximizing power, can depend crucially on the nature of the of the unknown distributions under investigation. The median performs best for the situation in Figure 5**C**, but situations are encountered where it trims too much, given the goal of minimizing the standard error. Roughly, a 20% trimmed mean competes reasonably well with the mean under normality. But as we move toward distributions that are more likely to generate outliers, at some point the median will have a smaller standard error than a 20% trimmed mean. Illustrations in section 5 demonstrate that this is a practical concern.

It is not being suggested that the mere presence of outliers will necessarily result in higher power when using a 20% trimmed mean or median. But it is being argued that simply ignoring the potential impact of outliers can be a serious practical concern.

In terms of controlling the Type I error probability, effective techniques are available for both the 20% trimmed mean and median. Because the choice between a 20% trimmed mean and median is not straightforward in terms of maximizing power, it is suggested that in exploratory studies, both of these estimators be considered.

When dealing with least squares regression or Pearson’s correlation, again outliers are a serious concern. Indeed, even a single outlier might give a highly distorted sense about the association among the bulk of the participants under study. In particular, important associations might be missed. One of the more obvious ways of dealing with this issue is to switch to Kendall’s tau or Spearman’s rho. However, these measures of associations do not deal with all possible concerns related to outliers. For instance, two outliers, properly placed, can give a distorted sense about the association among the bulk of the data (e.g, Wilcox, 2017b, p. 239).

A measure of association that deals with this issue is the skipped correlation where outliers are detected using a projection method, these points are removed, and Pearson’s correlation is computed using the remaining data. Complete details are summarized in Wilcox (2017a). This particular skipped correlation can be computed with the R function scor and a confidence interval, that allows heteroscedasticity, can be computed with scorci. This function also reports a p-value when testing the hypothesis that the correlation is equal to zero. (See section 3.2 for a description of common mistakes when testing hypotheses and outliers are removed.) Matlab code is available too (Pernet, Wilcox, & Rousselet, 2012).

### 2.4 Curvature

Typically, a regression line is assumed to be straight. In some situations, this approach seems to suffice. However, it cannot be stressed too strongly that there is a substantial literature indicating that this is not always the case. A vast array of new and improved nonparametric methods for dealing with curvature is now available, but complete details go beyond the scope of this paper. Here it is merely remarked that among the many nonparametric regression estimators that have been proposed, generally known as smoothers, two that seem to be particularly useful are Cleveland’s (1979) estimator, which can be applied via the R function lplot, and the running-interval smoother (Wilcox, 2017a), which can be applied with the R function rplot. Arguments can be made that other smoothers should be given serious consideration. Readers interested in these details are referred to Wilcox (2017a, c).

Cleveland’s smoother was initially designed to estimate the mean of the dependent variable given some value of the independent variable. The R function contains an option for dealing with outliers among the dependent variable, but it currently seems that the running-interval smoother is generally better for dealing with this issue. By default, the running-interval smoother estimates the 20% trimmed mean of the dependent variable, but any other measure of central tendency can be used via the argument est.

A simple strategy for dealing with curvature is to include a quadratic term. Let *X* denote the independent variable. An even more general strategy is to include *X^a^* in the model for some appropriate choice for the exponent *a*. But this strategy can be unsatisfactory as illustrated in section 5.4 using data from a study dealing with fractional anisotropy and reading ability. In general, smoothers provide a more satisfactory approach.

There are methods for testing the hypothesis that a regression line is straight (e.g., Wilcox, 2917a). However, failing to reject does not provide compelling evidence that it is safe to assume that indeed the regression line is straight. It is unclear when this approach has enough power to detect situations where curvature is an important practical concern. The best advice is to plot an estimate of the regression line using a smoother. If there is any indication that curvature might be an issue, use modern methods for dealing with curvature, many of which are summarized in Wilcox (2017a, c).

## 3 Dealing with Violation of Assumptions

Based on conventional training, there are some seemingly obvious strategies for dealing with the concerns reviewed in the previous section. But by modern standards, generally these strategies are relatively ineffective. This section summarizes strategies that perform poorly, followed by a brief description of modern methods that give improved results.

### 3.1 Testing Assumptions

A seemingly natural strategy is to test assumptions. In particular, test the hypothesis that distributions have a normal distribution and test the hypothesis that there is homoscedasticity. This approach generally fails, roughly because such tests do not have enough power to detect situations where violating these assumptions is a practical concern. That is, these tests can fail to detect situations that have an inordinately detrimental influence on statistical power and parameter estimation.

For example, Wilcox (2017a, c) lists six studies aimed at testing the homoscedasticity assumption with the goal of salvaging a method that assumes homoscedasticity (cf., Keselman et al., 2016). Briefly, these simulation studies generate data from a situation where it is known that homoscedastic methods perform poorly in terms of controlling the Type I error when there is heteroscedasticity. Then various methods for testing the homoscedasticity are performed, and if they reject, a heteroscedastic method is used instead. All six studies came to the same conclusion: this strategy is unsatisfactory. Presumably there are situations where this strategy is satisfactory, but it is unknown how to accurately determine whether this is the case based on the available data. In practical terms, all indications are that it is best to always use a heteroscedastic method when comparing measures of central tendency, and when dealing with regression as well as measures of association such as Pearson’s correlation.

As for testing the normality assumption, for instance using the Kolmogorov–Smirnov test, currently a better strategy is to use a more modern method that performs about as well as conventional methods under normality, but which continues to perform relatively well in situations where standard techniques perform poorly. There are many such methods (e.g., Wilcox, 2017a, c), some of which are outlined in section 4. These methods can be applied using extant software as will be illustrated.

### 3.2 Outliers: Two Common Mistakes

There are two common mistakes regarding how to deal with outliers. The first is to search for outliers using the mean and standard deviation. For example, declare the value *X* and outlier if
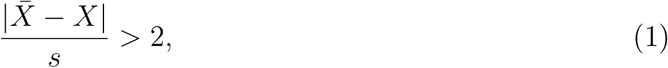

where 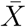 and *s* are the usual sample mean and standard deviation, respectively. A problem with this strategy is that it suffers from masking, simply meaning that the very presence of outliers causes them to be missed (e.g, Rousseeuw & Leroy, 1987; Wilcox, 2017a).

Consider, for example, the values 1, 2, 2, 3, 4, 6, 100 and 100. The two last observations appear to be clear outliers, yet the rule given by (1) fails to flag them as such. The reason is simple: the standard deviation of the sample is very large, at almost 45, because it is not robust to outliers.

This is not to suggest that all outliers will be missed; this is not necessarily the case. The point is that multiple outliers might be missed that adversely affect any conventional method that might be used to compare means. Much more effective are the boxplot rule and the so-called MAD-median rule.

The boxplot rule is applied as follows. Let *q*_1_ and *q*_2_ be estimates of the lower and upper quartiles, respectively. Then the value *X* is declared an outlier if *X* < *q*_1_ – 1.5(*q*_2_ − *q*_1_) or if *X* > *q*_2_ + 1.5(*q*_2_ − *q*_1_) As for the MAD-median rule, let *X*_1_,…, *X_n_* denote a random sample and let *M* be the usual sample median. MAD (the median absolute deviation to the median) is the median of the values |*X*_1_ – *M*|, … ,|*X_a_* – *M*|. The MAD-median rule declares the value X an outlier if 
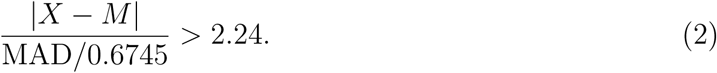

Under normality, it can be shown that MAD /0.6745 estimates the standard deviation, and of course *M* estimates the population mean. So the MAD-median rule is similar to using (1), only rather than use a two-standard deviation rule, 2.24 is used instead.

As an illustration, consider the values 1.85, 1.11, 1.11, 0.37, 0.37, 1.85, 71.53, and 71.53. The MAD-median rule detects the outliers: 71.53. But the rule given by (1) does not. The MAD-median rule is better than the boxplot rule in terms of avoiding masking.

The second mistake is discarding outliers and applying some standard method for comparing means using the remaining data. This results in an incorrect estimate of the standard error, regardless how large the sample size might be. That is, an invalid test statistic is being used. Roughly, it can be shown that the remaining data are dependent, they are correlated, which invalidates the derivation of the standard error. Of course, if an argument can be made that a value is invalid, discarding it is reasonable and does not lead to technical issues. For instance, a straightforward case can be made if a measurement is outside physiological bounds, or if it follows a biologically non-plausible pattern over time, such as during an electrophysiological recording. But otherwise, the estimate of the standard error can be off by a factor of 2 (e.g., Wilcox, 2017b), which is a serious practical issue. A simple way of dealing with this issue, when using a 20% trimmed mean or median, is to use a percentile bootstrap method. (With reasonably large sample sizes, alternatives to the percentile bootstrap method can be used, which are described in Wilcox 2017a, c). The main point here is that these methods are readily applied with the free software R, which is playing an increasing role in basic training. Some illustrations are given in section 5.

It is noted that when dealing with regression, outliers among the independent variables can be removed when testing hypotheses. But if outliers among the dependent variable are removed, conventional hypothesis testing techniques based on the least squares estimator are no longer valid, even when there is homoscedasticity. Again, the issue is that an incorrect estimate of the standard error is being used. When using robust regression estimators that deal with outliers among the dependent variable, again a percentile bootstrap method can be used to test hypotheses. Complete details are summarized in Wilcox (2017a, c). There are numerous regression estimators that effectively deal with outliers among the dependent variable, but a brief summary of the many details is impossible. The Theil and Sen estimator as well as the MM-estimator are relatively good choices, but arguments can be made that alternative estimators deserve serious consideration.

### 3.3 Transform the Data

A common strategy for dealing with non-normality or heteroscedasticity is to transform the data. There are exceptions, but generally this approach is unsatisfactory for several reasons. First, the transformed data can again be skewed to the point that classic techniques perform poorly (e.g., Wilcox, 2017b). Second, this simple approach does not deal with outliers in a satisfactory manner (e.g., Doksum & Wong, 1983; Rasmussen, 1989). The number of outliers might decline, but it can remain the same and even increase. Currently, a much more satisfactory strategy is to use a modern robust method such as a bootstrap method in conjunction with a 20% trimmed mean or median. This approach also deals with heteroscedasticity in a very effective manner (e.g., Wilcox, 2017a, c).

Another concern is that a transformation changes the hypothesis being tested. In effect, transformations muddy the interpretation of any comparison because a transformation of the data also transforms the construct that it measures (Grayson, 2004).

### 3.4 Use a Rank-Based Method

Standard training suggests a simple way of dealing with non-normality: use a rank-based method such as the WMW test, the Kruskal–Wallis test and Friedman’s method. The first thing to stress is that under general conditions these methods are not designed to compare medians or other measures of central tendency. (For an illustration based on the Wilcoxon signed rank test, see Wilcox, 2017b, p. 367. Also see Fagerland & Sandvik, 2009.) Moreover, the derivation of these methods is based on the assumption that the groups have identical distributions. So in particular, homoscedasticity is assumed. In practical terms, if they reject, conclude that the distributions differ.

But to get a more detailed understanding of how groups differ and by how much, alternative inferential techniques should be used in conjunction with plots such as those summarized by Rousselet, et al. (2017). For example, use methods based on a trimmed mean or median.

Many improved rank-based methods have been derived (Brunner et al., 2002). But again, these methods are aimed at testing the hypothesis that groups have identical distributions. Important exceptions are the improvements on the WMW test (e.g., Cliff, 1996; Wilcox, 2017a, b), which, as previously noted, are aimed at making inferences about the probability that a random observation from the first group is less than a random observation from the second.

### 3.5 Permutation Methods

For completeness, it is noted that permutation methods have received some attention in the neuroscience literature (e.g., Winkler et al., 2014; Pernet et al., 2015). Briefly, this approach is well designed to test the hypothesis that two groups have identical distributions. But based on results reported by Boik (1987), this approach cannot be recommended when comparing means. The same is true when comparing medians for reasons summarized by Romano (1990). Chung and Romano (2013) summarize general theoretical concerns and limitations. They go on to suggest a modification of the standard permutation method, but at least in some situations the method is unsatisfactory (Wilcox, 2017c, section 7.7). A deep understanding of when this modification performs well is in need of further study.

### 3.6 More Comments about the Median

In terms of power, the mean is preferable over the median or 20% trimmed mean when dealing with symmetric distributions for which outliers are rare. If the distribution are skewed, the median and 20% trimmed mean can better reflect what is typical, and improved control over the Type I error probability can be achieved. When outliers occur, there is the possibility that the mean will have a much larger standard error than the median or 20% trimmed mean. Consequently, methods based on the mean might have relatively poor power. Note, however, that for skewed distributions, the difference between two means might be larger than the difference between the corresponding medians. Consequently, even when outliers are common, it is possible that a method based on the means will have more power. In terms of maximizing power, a crude rule is to use a 20% trimmed mean, but the seemingly more important point is that no method dominates. Focusing on a single measure of central tendency might result is missing an important difference. So again, exploratory studies can be vitally important.

Even when there are tied values, it is now possible to get excellent control over the probability of a Type I error when using the usual sample median. For the one-sample case, this can be done with using a distribution free technique via the R function sintv2. Distribution free means that the actual Type I error probability can be determined exactly assuming random sampling only. When comparing two or more groups, currently the only known technique that performs well is the percentile bootstrap. Methods based on estimates of the standard error can perform poorly, even with large sample sizes. Also, when there are tied values, the distribution of the sample median does not necessarily converge to a normal distribution as the sample size increases. The very presence of tied values is not necessarily disastrous. But it is unclear how many tied values can be accommodated before disaster strikes. The percentile bootstrap method eliminates this concern.

## 4 Comparing Groups and Measures of Association

This section elaborates on methods aimed at comparing groups and measures of association. First attention is focused on two independent groups. Comparing dependent groups is discussed in section 4.4. Section 4.5 comments briefly on more complex designs. This is followed by a description of how to compare measures of association as well as an indication of modern advances related to the analysis of covariance. Included are indications of how to apply these methods using the free software R, which at the moment is easily the best software for applying modern methods. R is a vast and powerful software package. Certainly matlab could be used, but this would require writing hundreds of functions in order to compete with R. There are numerous books on R, but only a relatively small subset of the basic commands is needed to apply the functions described here. (See, for example, Wilcox, 2017b, c.)

The R functions noted here are stored in the R package WRS, which can be installed as indicated at https://github.com/nicebread/WRS. Alternatively, and seemingly easier, use the R command source on the file Rallfun-v33.txt, which can be downloaded from http://dornsife.usc.edu/labs/rwilcox/software/.

All recommended methods deal with heteroscedasticity. When comparing groups and distributions differ in shape, these methods are generally better than classic methods for comparing means, which can perform poorly.

### 4.1 Dealing with Small Sample Sizes

This section focuses on the common situation in the neuroscience where the sample sizes are relatively small. When the sample sizes are very small, say less than or equal ten and greater than four, conventional methods based on means are satisfactory in terms of Type I errors when the null hypothesis is that the groups have identical distributions. If the goal is to control the probability of a Type I error when the null hypothesis is that groups have equal means, extant methods can be unsatisfactory. And as previously noted, methods based on means can have poor power relative to alternative techniques.

Many of the more effective methods are based in part on the percentile bootstrap method. Consider, for example, the goal of comparing the medians of two independent groups. Let *M_x_* and *M_y_* be the sample medians for the two groups being compared and let *D* = *M_x_* –*M_y_.* The basic strategy is to perform a simulation based on the observed data with the goal of approximating the distribution of *D*, which can then be used to compute a p-value as well as a confidence interval.

Let *n* and *m* denote the sample sizes for the first and second group, respectively. The percentile bootstrap method proceeds as follows. For the first group, randomly sample with replacement *n* observations. This yields what is generally called a bootstrap sample. For the second group, randomly sample with replacement *m* observations. Next, based on these bootstrap samples, compute the sample medians, say 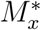 and 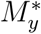. Let 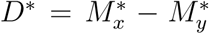. Repeating this process many times, a p-value (and confidence interval) can be computed based on the proportion of times *D*^*^ < 0 as well as the proportion of times *D*^*^ = 0. (More precise details are given in Wilcox, 2017a, c.) This method has been found to provide reasonably good control over the probability of a Type I error when both *n* and *m* are greater than equal to five. The R function medpb2 performs this technique.

The method just described performs very well compared to alternative techniques. In fact, regardless of how large the sample sizes might be, the percentile bootstrap method is the only known method that continues to perform reasonably well when comparing medians and there are duplicated values. (Also see Wilcox (2017a, section 5.3.)

For the special case where the goal is to compare means, there is no method that provides reasonably accurate control over the Type I error probability for a relatively broad range of situations. In fairness, situations can be created where means perform well and indeed have higher power than methods based on a 20% trimmed mean or median. When dealing with perfectly symmetric distributions where outliers are unlikely to occur, methods based on means, and that allow heteroscedasticity, perform relatively well. But with small sample sizes, there is no satisfactory diagnostic tool indicating whether distributions satisfy these two conditions in an adequate manner. Generally, using means comes with the relatively high risk of poor control over the Type I error probability and relatively poor power.

Switching to a 20% trimmed mean, the method derived by Yuen (1974) performs fairly well even when the smallest sample size is six (cf. Özdemir et al., 2013). It can be applied with the R function yuen. (Yuen’s method reduces to Welch’s method for comparing means when there is no trimming.) When the smallest sample size is five, it can be unsatisfactory in situations where the percentile bootstrap method, used in conjunction with the median, continues to perform reasonably well. A rough rule is that the ability to control the Type I error probability improves as the amount of trimming increases. With small sample sizes, and when the goal is to compare means, it is unknown how to control the Type I error probability reasonably well over a reasonably broad range of situations.

Another approach is to focus on *P*(*X* < *Y*), the probability that a randomly sample observation from the first group is less than a randomly sample observation from the second group. This strategy is based in part on an estimate of the distribution of *D* = *X* – *Y*, the distribution of all pairwise differences between observations in each group.

To illustrate this point, let *X*_1_, … , *X_n_* and *Y*_1_,… , *Y_m_* be random samples of size *n* and *m*, respectively. Let *D_ik_* = *X_i_* – *Y_k_* (i = 1, … *n*; *k* = 1, … , *m*). Then the usual estimate of *P*(*X* < Y) is simply the proportion of *D_ik_* values less than zero. For instance, if *X* = (1, 2, 3) and *Y* = (1, 2.5, 4), then *D* = (0.0, 1.0, 2.0, -1.5, -0.5, 0.5, -3.0, -2.0, -1.0) and the estimate of *P*(*X* < *Y*) is 4/9, the proportion of *D* values less than zero.

Let *μ_x_, μ_y_* and *_μD_* denote the population means associated with *X, Y* and *D*, respectively. From basic principles, *μ_x_* – *μ_y_* = *μ_D_.* That is, the difference between two means is the same as the mean of all pairwise differences. However, let *θ_x_*, *θ_y_* and *θ_D_* denote the population medians associated with *X*, *Y* and *D*, respectively. For symmetric distributions, *θ_x_* – *θ_y_* = *θ_D_*, but otherwise it is generally the case that *θ_x_* – *θ_y_* ≠ *θ_D_.* In other words, the difference between medians is typically not the same as the median of all pairwise differences. The same is true when using any amount of trimming greater than zero. Roughly, *θ_x_* and *θ_y_* reflect the typical response for each group, while *θ_D_* reflects the typical difference between two randomly sampled participants, one from each group. Although less known, the second perspective can be instructive in many situations. For instance, in a clinical setting in which we want to know what effect to expect when randomly sampling a patient and a control participant.

If two groups do not differ in any manner, *P*(*X* < *Y*) = 0.5. Consequently, a basic goal is testing
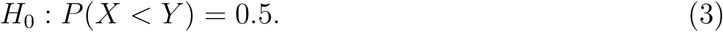

If this hypothesis is rejected, this indicates that it is reasonable to make a decision about whether *P*(*X* < *Y*) is less than or greater than 0.5 It is readily verified that this is the same as testing
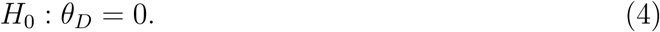

An appeal of *P*(*X* < *Y*) is that it is easily understood by non-statisticians, and it has practical importance for reasons summarized, among others, by Cliff (1996), Ruscio (2008) and Newcombe (2006). The Wilcoxon–Mann–Whitney (WMW) test is based on an estimate *P*(*X* < *Y*). However, it is unsatisfactory in terms of making inferences about this probability because the estimate of the standard error assumes that the distributions are identical. If the distributions differ, an incorrect estimate of the standard error is being used. More modern methods deal with this issue. The method derived by Cliff (1996) for testing (3) performs relatively well with small sample sizes and can be applied via the R function cidv2. (We are not aware of any commercial software package that contains this method.)

However, there is the complication that for skewed distributions, differences among the means, for example, can be substantially smaller as well as substantially larger than differences among 20% trimmed means or medians. That is, regardless of how large the sample sizes might be, power can be substantially impacted by which measure of central tendency is used.

### 4.2 Comparing Quantiles

Rather than compare groups based on a single measure of central tendency, typically the mean, another approach is to compare multiple quantiles. For example, compare the quartiles, or all of the deciles, or even all quantiles. This provides more detail about where and how the two distributions differ (Rousselet, Pernet & Wilcox, 2017). For example, the typical participants might not differ very much based on the medians, but the reverse might be true among low scoring individuals.

First consider the goal of comparing all quantiles in a manner that controls the probability of one or more Type I errors among all the tests that are performed. Assuming random sampling only, Doksum and Sievers (1976) derived such a method that can be applied via the R function sband. The method is based on a generalization of the Kolmogorov–Smirnov test. A negative feature is that power can be adversely affected when there are tied values. And when the goal is to compare the more extreme quantiles, again power might be relatively low. A way of reducing these concerns is to compare the deciles using a percentile bootstrap method in conjunction with the quantile estimator derived by Harrell and Davis (1982). This is easily done with the R functions qcomhd.

Note that if the distributions associated with *X* and *Y* do not differ, then *D* = *X* − *Y* will have a symmetric distribution about zero. Let *x_q_* be the qth quantile of *D*, 0 < *q* < 0.5. In particular, it will be the case that *x_q_* + *x*_1−*q*_ = 0 when *X* and *Y* have identical distributions. The median (2nd quartile) will be zero, and, for instance, the sum of the 3rd quartile (0.75 quantile) and the 1st quartile (0.25 quantile) will be zero. So this sum provides yet another perspective on how distributions differ (see illustrations in Rousselet, Pernet & Wilcox, 2017).

Imagine, for example, that an experimental group is compared to a control group based on some measure of depressive symptoms. If *x*_0.25_ = −4 and *x*_0.75_ = 6, then for a single randomly sampled observation from each group, there is a sense in which the experimental treatment outweighs no treatment, because positive differences (beneficial effect) tend to be larger than negative differences (detrimental effect). The hypothesis 
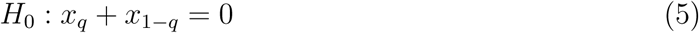
can be tested with the R function cbmhd. A confidence interval is returned as well. Current results indicate that the method provides reasonably accurate control over the Type I error probability when *q* = 0.25 and the sample sizes are greater than or equal to ten. For *q* = 0.1, sample sizes greater than or equal to twenty should be used (Wilcox, 2012).

### 4.3 Eliminate Outliers and Average the Remaining Values

Rather than use means, trimmed means or the median, another approach is to use an estimator that down weights or eliminates outliers. For example, use the MAD-median to search for outliers, remove any that are found and average the remaining values. This is generally known as a modified one-step M-estimator (MOM). This approach might seem preferable to using a trimmed mean or median because trimming can eliminate points that are not outliers. But this issue is far from simple. Indeed, there are indications that when testing hypotheses, the expectation is that using a trimmed mean or median will perform better in terms of Type I errors and power (Wilcox, 2017a). However, there are exceptions: no single estimator dominates.

As previously noted, an invalid strategy is to eliminate extreme values and apply conventional methods for means based on the remaining data, because the wrong standard error is used. Switching to a percentile bootstrap deals with this issue when using MOM as well as related estimators. The R function pb2gen applies this method.

### 4.4 Comparing Dependent Variables

Next, consider the goal of comparing two dependent variables. That is, the variables might be correlated. Based on the random sample (*X*_1_, *Y*_1_), … , (*X_n_*, *Y_n_*), let *D_i_* = *X_i_* – *Y_i_* (*i* = 1, … , *n*). Even when *X* and *Y* are correlated, *μ_x_* – *μ_y_* = *μ_D_*, the difference between the population means is equal to the mean of the difference scores. But under general conditions this is not the case when working with trimmed means. When dealing with medians, for example, it is generally the case that *θ_x_* – *θ_y_* ≠ *θ_D_.*

If the distribution of *D* is symmetric and light-tailed (outliers are relatively rare), the paired t-test performs reasonably well. As we move toward a skewed distribution, at some point this is no longer the case for reasons summarized in section 2.1. Moreover, power and control over the probability of a Type I error are also a function of the likelihood of encountering outliers.

There is a method for computing a confidence interval for *θ_D_* for which the probability of a Type I error can be determined exactly assuming random sampling only (e.g., Hettmansperger & McKean, 1998). In practice, a slight modification of this method is recommended that was derived by Hettmansperger and Sheather (1986). So when sample sizes are very small, this method performs very well in terms of controlling the probability of a Type I error. And in general, it is an excellent method for making inferences about *θ_D_.* The method can be applied via the R function sintv2.

As for trimmed means, with the focus still on *D*, a percentile bootstrap method can be used via the R function trimpb or wmcppb. Again, with 20% trimming, reasonably good control over the Type I error probability can be achieved. With *n* = 20, the percentile bootstrap method is better than the non-bootstrap method derived by Tukey and McLaughlin (1963). With large enough sample sizes the Tukey–McLaughlin method can be used in lieu of the percentile bootstrap method via the R function trimci, but it is unclear just how large the sample size must be.

In some situations, there might be interest in comparing measures of central tendencies associated with the marginal distributions rather than the difference scores. Imagine, for example, participants consist of married couples. One issue might be the typical difference between a husband and his wife, in which case difference scores would be used. Another issue might be how the typical male compares to the typical female. So now the goal would be to test *H*_0_: *θ_x_* = *θ_y_*, rather than *H*_0_: *θ_D_* = 0. The R function dmeppb tests the first of these hypotheses and performs relatively well, even when there are tied values. If the goal is to compare the marginal trimmed means, rather than make inferences about the trimmed mean of the difference scores, use the R function dtrimpb, or use wmcppb and set the argument dif=FALSE. When dealing with a moderately large sample size, the R function yuend can be used instead, but there is no clear indication just how large the sample size must be. A collection of quantiles can be compared with Dqcomhd and all of the quantiles can be compared via the function lband.

Yet another approach is to use the classic sign test, which is aimed at making inferences about *P*(*X* < *Y*). As is evident, this probability provides a useful perspective on the nature of the difference between the two dependent variables under study beyond simply comparing measures of central tendency. The R function signt performs the sign test, which by default uses the method derived by Agresti and Coull (1998). If the goal is to ensure that the confidence interval has probability coverage at least 1 – *α*, rather than approximately equal to 1 – α, the Schilling and Doi (2014) method can be used by setting the argument SD=TRUE when using the R function signt. In contrast to the Schilling and Doi method, p-values can be computed when using the Agresti and Coull technique. Another negative feature of the Schilling and Doi method is that execution time can be extremely high even with a moderately large sample size.

A criticism of the sign test is that its power might be lower than the Wilcoxon signed rank test. However, this issue is not straightforward. Moreover, the sign test can reject in situations where other conventional methods do not. Again, which method has the highest power depends on the characteristics of the unknown distributions generating the data. Also, in contrast to the sign test, the Wilcoxon signed rank test provides no insight into the nature of any difference that might exist without making rather restrictive assumptions about the underlying distributions. In particular, under general conditions, it does not compare medians or some other measure of central tendency as previously noted.

### 4.5 More Complex Designs

It is noted that when dealing with a one-way or higher ANOVA design, violations of the normality and homoscedasticity assumptions, associated with classic methods for means, become an even more serious issue in terms of both Type I error probabilities and power. Robust methods have been derived (Wilcox, 2017a, c), but the many details go beyond the scope of this paper. However, a few points are worth stressing.

Momentarily assume normality and homoscedasticity. Another important insight has to do with the role of the ANOVA F test versus post-hoc multiple comparison procedures such as the Tukey–Kramer method. In terms of controlling the probability of one or more Type I errors, is it necessary to first reject with the ANOVA F test? The answer is an unequivocal no. With equal sample sizes, the Tukey–Kramer method provides exact control. But if it is used only after the ANOVA F test rejects, this is no longer the case; it is lower than the nominal level (Bernhardson, 1975). For unequal sample sizes, the probability of one or more Type I errors is less than or equal to the nominal level when using the Tukey–Kramer method. But if it is used only after the ANOVA F test rejects, it is even lower, which can negatively impact power. More generally, if an experiment aims to test specific hypotheses involving subsets of conditions, there is no obligation to first perform an ANOVA: the analyses should focus directly on the comparisons of interest using, for instance, the functions for linear contrasts listed below.

Now consider non-normality and heteroscedasticity. When performing all pairwise comparisons, for example, most modern methods are designed to control the the probability of one or more Type I errors without first performing a robust analog of the ANOVA F test. There are, however, situations where a robust analog of the ANOVA F test can help increase power (e.g., Wilcox, 2017a, section 7.4).

For a one-way design where the goal is to compare all pairs of groups, a percentile bootstrap method can be used via the R function linconpb. A non-bootstrap method is performed by lincon. For medians, use medpb. For an extension of Cliff’s method, use cidmul. Methods and corresponding R functions for both two-way and three-way designs, including techniques for dependent groups, are available as well; see Wilcox (2017a, c).

### 4.6 Comparing Independent Correlations and Regression Slopes

Next, consider two independent groups where for each group there is interest in the strength of the association between two variables. A common goal is to test the hypothesis that the strength of association is the same for both groups.

Let *ρ_j_* (*j* = 1, 2) be Pearson’s correlation for the *j*th group and consider the goal of testing
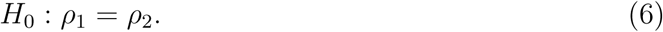

Various methods for accomplishing this goal are known to be unsatisfactory (Wilcox, 2009). For example, one might use Fisher’s r-to-z transformation, but it follows immediately from results in Duncan and Layard (1973) that this approach performs poorly under general conditions. Methods that assume homoscedasticity, as depicted in Figure 4, can be unsatisfactory as well. As previously noted, when there is an association (the variables are dependent), and in particular there is heteroscedasticity, a practical concern is that the wrong standard error is being used when testing hypotheses about the slopes. That is, the derivation of the test statistic is valid when there is no association; independence implies homoscedasticity. But under general conditions it is invalid when there is heteroscedasticity. This concern extends to inferences made about Pearson’s correlation.

There are two methods that perform relatively well in terms of controlling the Type I error probability. The first is based on a modification of the basic percentile bootstrap method. Imagine that (6) is rejected if the confidence interval for *ρ*_1_ – *ρ*_2_ does not contain zero. So the Type I error probability depends on the width of this confidence interval. If it is too short, the actual Type I error probability will exceed 0.05. With small sample sizes this is exactly what happens when using the basic percentile bootstrap method. The modification consists of widening the confidence interval for *ρ*_1_ – *ρ*_2_ when the sample size is small. The amount it is widened depends on the sample sizes. The method can be applied via the R function twopcor. A limitation is that this method can be used only when the Type I error is 0.05 and it does not yield a p-value.

The second approach is to use a method that estimates the standard error in a manner that deals with heteroscedasticity. When dealing with the slope of the least squares regression line, several methods are now available for getting valid estimates of the standard error when there is heteroscedasticity (Wilcox, 2017a). One of these is called the HC4 estimator, which can be used to test (6) via the R function twohc4cor.

As previously noted, Pearson’s correlation is not robust: even a single outlier might substantially impact its value giving a distorted sense of the strength of the association among the bulk of the points. Switching to Kendall’s tau or Spearman’s rho, now a basic percentile bootstrap method can be used to compare two independent groups, in a manner that allows heteroscedasticity, via the R function twocor. As noted in section 2.3, the skipped correlation can be used via the R function scorci.

The slopes of regression lines can be compared as well using methods that allow heteroscedasticity. For least squares regression, use the R function ols2ci. For robust regression estimators, use reg2ci.

### 4.7 Comparing Correlations, the Overlapping Case

Now consider a single dependent variable *Y* and two independent variables, *X*_1_ and *X*_2_. A common and fundamental goal is understanding the relative importance of *X*_1_ and *X*_2_ in terms of their association with *Y.* A typical mistake in neuroscience is to perform two separate tests of associations, one between *X*_1_ and *Y*, another between *X*_2_ and *Y*, without explicitly comparing the association strengths between the independent variables (Nieuwenhuis et al. 2011). For instance, reporting that one test is significant, and the other is not, cannot be used to conclude that the associations themselves differ. A common example would be when an association is estimated between each of two brain measurements and a behavioural outcome.

Their are many methods for estimating which independent variable is more important, many of which are known to be unsatisfactory (e.g., Wilcox, 2017c, section 6.13). Stepwise regression is among the unsatisfactory techniques for reasons summarized by Montgomery and Peck (1992, Section 7.2.3) as well as Derksen and Keselman (1992). Regardless of their relative merits, a practical limitation is that they do not reflect the strength of empirical evidence that the most important independent variable has been chosen. One strategy is to test *H*_0_: *ρ*_1_ = *ρ*_2_, where now *ρ_j_* (*j* = 1, 2) is the correlation between *Y* and *X_j_.* This can be done with the R function TWOpov. When dealing with robust correlations, use the function twoDcorR.

However, a criticism of this approach is that it does not take into account the nature of the association when both independent variables are included in the model. This is a concern because the strength of the association between *Y* and *X*_1_ can depend on whether *X*_2_ is included in the model as illustrated in section 5.4. There is now a robust method for testing the hypothesis that there is no difference in the association strength when both *X*_1_ and *X*_2_ are included in the model. (Wilcox, 2017a, 2016). Heteroscedasticity is allowed. If, for example, there are three independent variables, one can test the hypothesis that the strength of the association for the first two independent variables is equal to the strength of the association for the third independent variable. The method can be applied with the R function regIVcom. A modification and extension of the method has been derived when there is curvature (Wilcox, in press), but it is limited to two independent variables.

### 4.8 ANCOVA

The simplest version of the analysis of covariance (ANCOVA) consists of comparing the regression lines associated with two independent groups when there is a single independent variable. The classic method makes several restrictive assumptions: the regression lines are parallel, for each regression line there is homoscedasticity, the variance of the dependent variable is the same for both groups, normality, and a straight regression line provides an adequate approximation of the true association. Violating any of these assumptions is a serious practical concern. Violating two or more of these assumptions makes matters worse. There is now vast array of more modern methods that deal with violations of all of these assumptions Wilcox (2017a, chapter 12). These newer techniques can substantially increase power compared to the classic ANCOVA technique, and perhaps more importantly they can provide a deeper and more accurate understanding of how the groups compare. But the many details go beyond the scope of this paper.

As noted in the introduction, curvature is a more serious concern than is generally recognized. One strategy, as a partial check on the presence of curvature, is to simply plot the regression lines associated with two groups using the R functions lplot2g and rplot2g. When using these functions, as well as related functions, it can be vitally important to check on the impact of removing outliers among the independent variables. This is easily done with functions mentioned here by setting the argument xout=TRUE. If these plots suggest that curvature might be an issue, consider the R functions ancova and ancdet. This latter function applies method TAP in Wilcox (2017a, section 12.2.4) and can provide more detailed information about where and how two regression lines differ compared to the function ancova. These functions are based on methods that allow heteroscedasticity, non-normality, and they eliminate the classic assumption that the regression lines are parallel. For two independent variables, see Wilcox (2017a, section 12.4). If there is evidence that curvature is not an issue, again there are very effective methods that allow heteroscedasticity as well as non-normality (Wilcox, 2017a, section 12.1).

## 5 Some Illustrations

Using data from several studies, this section illustrates modern methods and how they contrast. Extant results suggest that robust methods have a relatively high likelihood of maximizing power, but as previously stressed, no single method dominates in terms of maximizing power. Another goal in this section is to underscore the suggestion that multiple perspectives can be important. More complete descriptions of the results, as well as the R code that was used, are available on figshare (Wilcox & Rousselet, 2017). The figshare reproducibility package also contains a more systematic assessment of type I error and power in the one-sample case. (See notebook `power_onesample.pdf`).

### 5.1 Spatial Acuity for Pain

The first illustration stems from Mancini et al. (2014) who report results aimed at providing a whole-body mapping of spatial acuity for pain. (Also see Mancini, 2016.) Here the focus is on their second experiment. Briefly, spatial acuity was assessed by measuring 2-point discrimination (2PD) thresholds for both pain and touch in 11 body territories. One goal was to compare touch measures taken at different body parts: forehead, shoulder, forearm, hand, back and thigh. Plots of the data are shown in Figure 6A for the six body parts. The sample size is *n* = 10.

**Figure 6:**
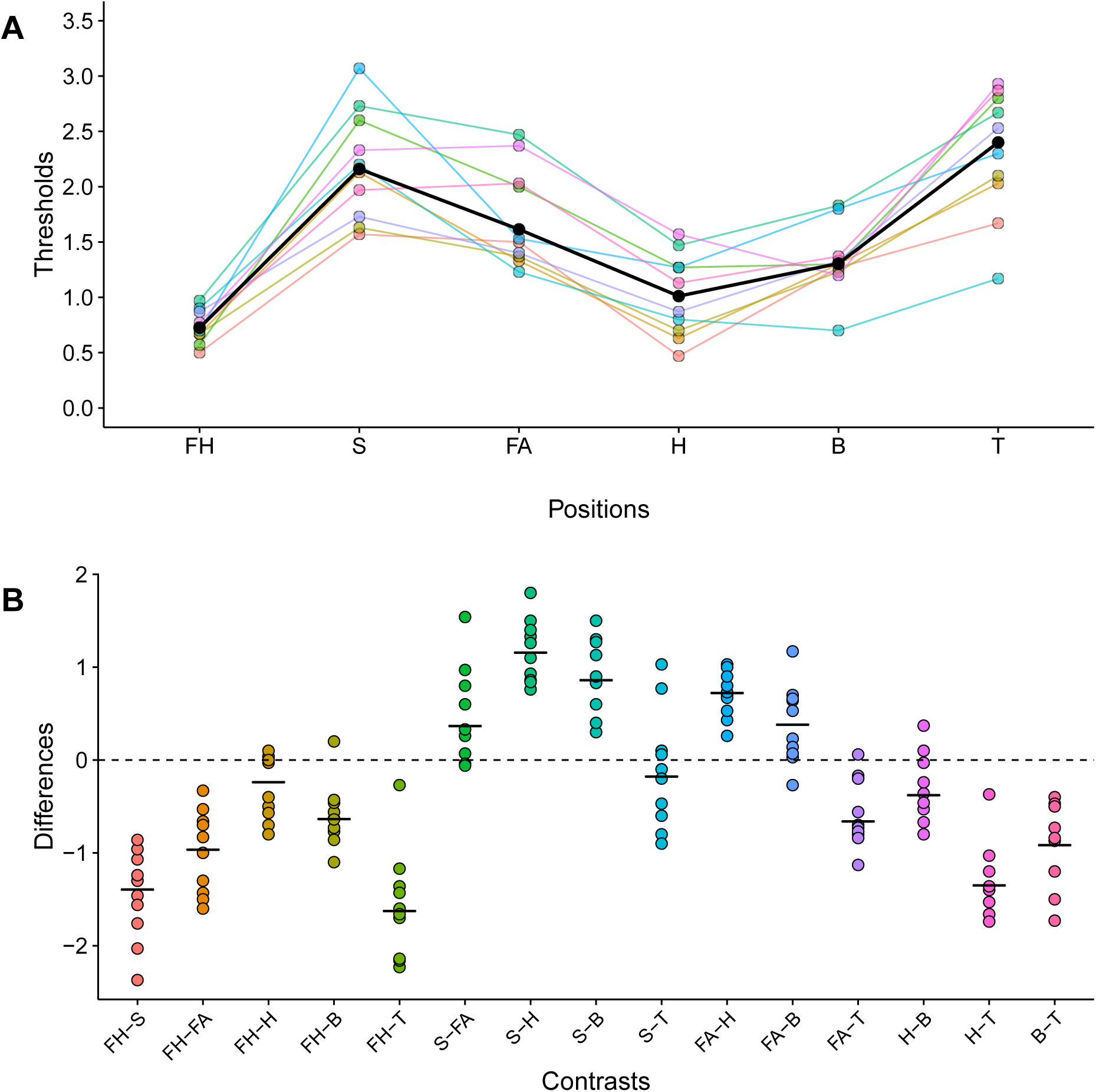
Data from Mancini et al. 2014. Panel A shows the marginal distributions of thresholds at locations FH = forehead, S = shoulder, FA = forearm, H = hand, B = back and T = thigh. Individual participants (n = 10) are shown as colored disks and lines. The medians across participants are shown in black. Panel B shows the distributions of all pairwise differences between the conditions shown in A. In each stripchart (1D scatterplot), the black horizontal line marks the median of the differences.

Their analyses were based on the ANOVA F test, followed by paired t-tests when the ANOVA F test was significant. Their significant results indicate that the distributions differ, but because the ANOVA F test is not a robust method when comparing means, there is some doubt about the nature of the differences. So one goal is to determine in which situations robust methods give similar results. And for the non-significant results, there is the issue of whether an important difference was missed due to using the ANOVA F test and Student’s t-test.

First we describe a situation where robust methods based a median and a 20% trimmed mean give reasonably similar results. Comparing foot and thigh pain measures based on Student’s t-test, the 0.95 confidence interval = [−0.112, 0.894] and the p-value is 0.112. For a 20% trimmed mean the 0.95 confidence interval is [−0.032, 0.778] and the p-value is 0.096. As for the median, the corresponding results were [−0.085, 0.75] and 0.062.

Next, all pairwise comparisons, based on touch, were performed for the following body parts: forehead, shoulder, forearm, hand, back and thigh. Figure 6B shows a plot of the difference scores for each pair of body parts. The probability of one or more Type I errors was controlled using an improvement on the Bonferroni method derived by Hochberg (1988). The simple strategy of using paired t-tests if the ANOVA F rejects does not control the probability of one or more Type I errors. If paired t-tests are used without controlling the probability of one or more Type I errors, as done by Mancini et al., 14 of the 15 hypotheses are rejected. If the probability of one or more Type I errors is controlled using Hochberg’s method, the following results were obtained. Comparing means via the R function wmcp (and the argument tr=0), 10 of the 15 hypotheses were rejected. Using medians via the R function dmedpb, 11 were rejected. As for the 20% trimmed mean, using the R function wmcppb, now 13 are rejected illustrating the point made earlier that the choice of method can make a practical difference.

It is noted that in various situations, using the sign test, the estimate of *P*(*X* < *Y*) was equal to one, which provides a useful perspective beyond using a mean or median.

### 5.2 Receptive Fields in Early Visual Cortex

The next illustrations are based on data analyzed by Talebi and Baker (2016) and presented in their Figure 9. The goal was to estimate visual receptive field (RF) models of neurones from a cat’s visual cortex using natural image stimuli. The authors provided a rich quantification of the neurones’ responses and demonstrated the existence of three functionally distinct categories of simple cells. The total sample size is 212.

There are three measures: latency, duration, and a direction selectivity index (dsi). For each of these measures there are three independent categories: nonoriented (nonOri) cells (n=101), expansive oriented (expOri) cells (n=48) and compressive oriented (compOri) cells (n=63).

First focus on latency. Talebi et al. used Student’s t-test to compare means. Comparing the means for nonOri and expOri, no significant difference is found. But an issue is whether Student’s t-test might be missing an important difference. The plot of the distributions shown in the top row of Figure 7**A**, which was created with the R function g5plot, provides a partial check on this possibility. As is evident, the distributions for nonOri and expOri are very similar suggesting that no method will yield a significant result. Using error bars is less convincing because important differences might exist when focusing instead on the median, 20% trimmed mean, or some other aspect of the distributions.

**Figure 7:**
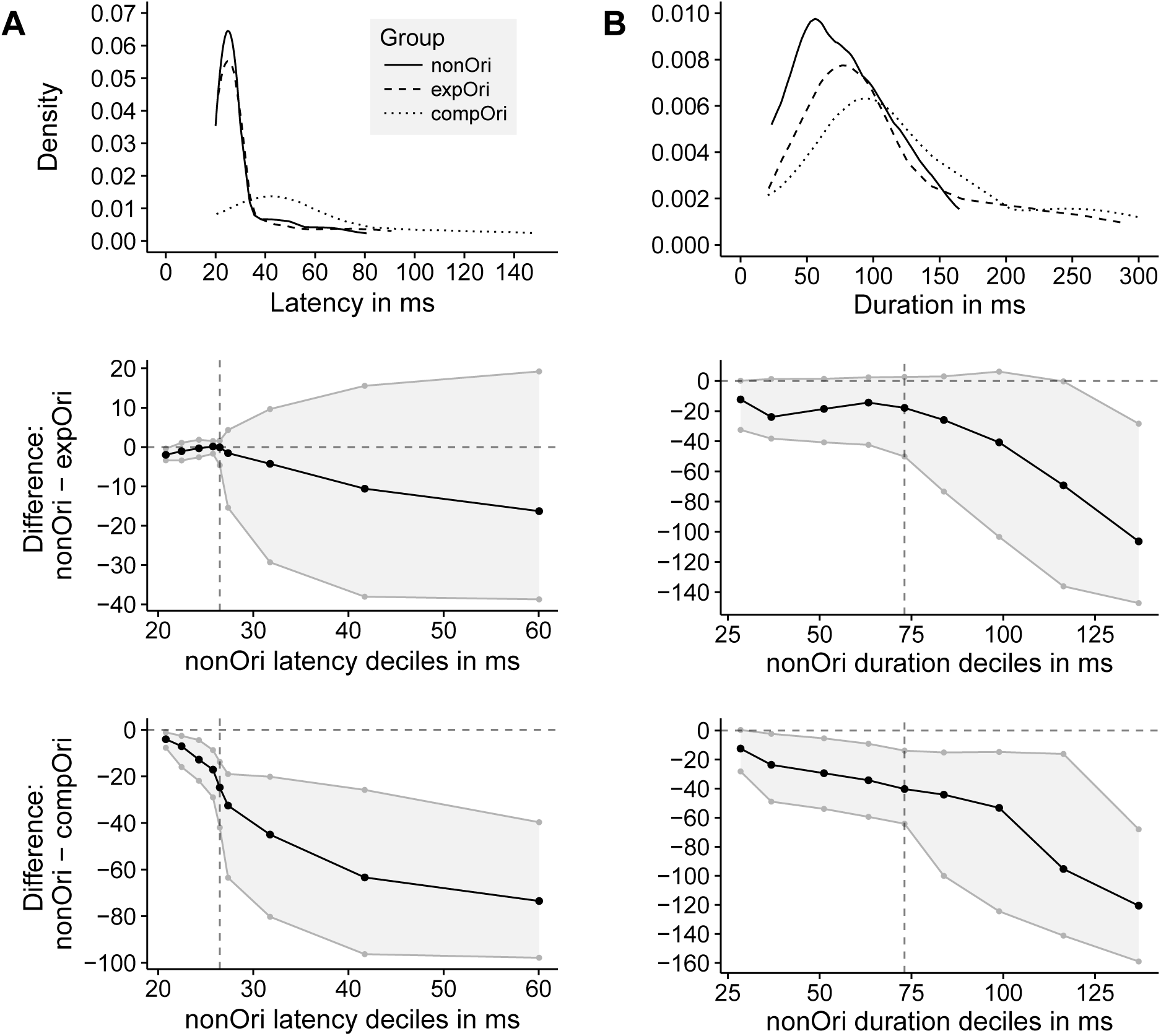
Response latencies A and durations B from cells recorded by Talebi and Baker (2016). Row 1: Estimated distributions of latency and duration measures for three categories: nonOri, expOri, and compOri. Rows 2 and 3 show shift functions based on comparisons between the deciles of two groups. The deciles of one group are on the x-axis; the difference between the deciles of the two groups is on the y-axis. The black dotted line is the difference between deciles, plotted as function of the deciles in one group. The grey dotted lines mark the 95% bootstrap confidence interval, which is also highlighted by the grey shaded area. Row 2: Differences between deciles for nonOri versus expOri, plotted as a function of the deciles for nonOri. Row 3: Differences between deciles for nonOri versus compOri, plotted as a function of the deciles for nonOri.

As noted in section 4.2, comparing the deciles can provide a more detailed understanding of how groups differ. The second row of Figure 7 shows the estimated difference between the deciles for the nonOri group versus the expOri group. The vertical dotted line indicates the median. Also shown are confidence intervals (computed via the R function qcomhd) having, approximately, simultaneous probability coverage equal to 0.95. That is, the probability of one or more Type I errors is approximately 0.05. In this particular case, differences between the deciles are more pronounced as we move from the lower to the upper deciles, but again no significant differences are found.

The third row shows the difference between the deciles for nonOri versus compOri. Again the magnitude of the differences becomes more pronounced moving from low to high measures. Now all of the deciles differ significantly except the 0.1 quantiles.

Next, consider durations in column B of Figure 7. Comparing nonOri to expOri using means, 20% trimmed means and medians, the corresponding p-values are 0.001, 0.005 and 0.009. Even when controlling the probability of one or more Type I errors using Hochberg’s method, all three reject at the 0.01 level. So a method based means rejects at the 0.01 level, but this merely indicates that the distributions differ in some manner. To provide an indication that the groups differ in terms of some measure of central tendency, using 20% trimmed means and medians is more satisfactory. The plot in row 2 of Figure 7B confirms an overall shift between the two distributions, and suggests a more specific pattern, with increasing differences in the deciles beyond the median.

Comparing nonOri to compOri, significant results were again obtained using means, 20% trimmed means and medians, the largest p-value is p=0.005. In contrast, qcomhd indicates a significant difference for all of the deciles excluding the 0.1 quantile. As can be seen from the last row in Figure 7B, once more the magnitude of the differences between the deciles increases as we move from the lower to the upper deciles. Again, this function provides a more detailed understanding of where and how the distributions differ significantly.

### 5.3 Mild Traumatic Brain Injury

The illustrations in this section stem from a study dealing with mild traumatic brain injury (Almeida-Suhett et al., 2014). Briefly, 5-6 week old male, Sprague–Dawley rats received a mild controlled cortical impact (CCI) injury. The dependent variable used here is the stereologically estimated total number of GAD-67-positive cells in the basolateral amygdala (BLA). Measures 1 and 7 days after surgery were compared to the sham-treated control group that received a craniotomy, but no CCI injury. A portion of their analyses focused on the ipsilateral sides of the BLA. Boxplots of the data are shown in Figure 8. The sample sizes are 13, 9 and 8, respectively.

**Figure 8:**
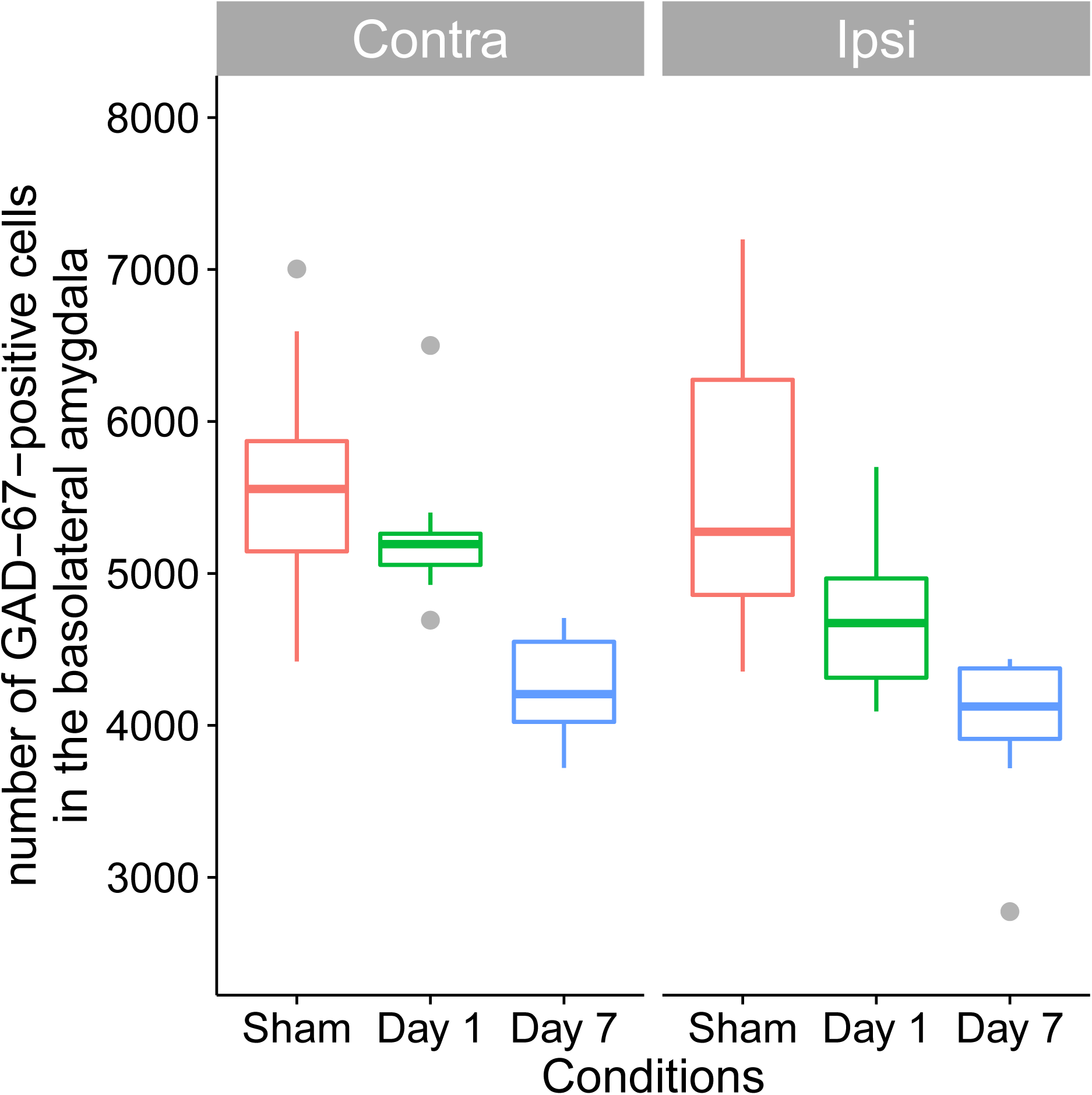
Boxplots for the contralateral and ipsilateral sides of the BLA

Almeida-Suhett et al. compared means using an ANOVA F test followed by Bonferroni post-hoc test. Comparing both day 1 and day 7 measures to the sham group based on Student’s t-test, the p-values are 0.0375 and *<* 0.001, respectively. So, if the Bonferroni method is used, the day 1 group does not differ significantly from the sham group when testing at the 0.05 level. However, using Hochberg’s improvement on the Bonferroni method, now the reverse decision is made.

Here, the day 1 group was compared again to the sham group based a percentile bootstrap method for comparing both 20% trimmed means and medians, as well as Cliff’s improvement on the WMW test. The corresponding p-values are 0.024, 0.086 and 0.040. If 20% trimmed means are compared instead with Yuen’s method, the p-value is p=0.079, but due to the relatively small sample sizes, a percentile bootstrap would be expected to provide more accurate control over the Type I error probability. The main point here is that the choice between Yuen and a percentile bootstrap method can make a practical difference. The boxplots suggest that sampling is from distributions that are unlikely to generate outliers, in which case a method based on the usual sample median might have relatively low power. When outliers are rare, a way of comparing medians that might have more power is to use instead the Harrell–Davis estimator mentioned in section 4.2 in conjunction with a percentile bootstrap method. Now p=0.049. Also, testing (4) with the R function wmwpb, p=0.031 So focusing on *θ_D_*, the median of all pairwise differences, rather than the individual medians, can make a practical difference.

In summary, when comparing the sham group to the Day 1 group, all of the methods that perform relatively well when sample sizes are small, described in section 4.1, reject at the 0.05 level except the percentile bootstrap method based on the usual sample median. Taken as a whole, the results suggest that measures for the sham group are typically higher than measures based on day 1 group. As for the day 7 data, now all of the methods used for the day 1 data have p-values less than or equal to 0.002.

The same analyses were done using the contralateral sides of the BLA. Now the results were consistent with those based on means: none are significant for day 1. As for the day 7 measures, both conventional and robust methods indicate significant results.

### 5.4 Fractional Anisotropy and Reading Ability

The next illustrations are based on data dealing with reading skills and structural brain development (Houston et al., 2014). The general goal was to investigate maturational volume changes in brain reading regions and their association with performance on reading measures. The statistical methods used were not made explicit. Presumably they were least squares regression or Pearson’s correlation coupled with the usual Student’s t-tests. The ages of the participants ranged between 6 and 16. After eliminating missing values, the sample size is *n* = 53. (It is unclear how Houston et al. dealt with missing values.)

As previously indicated, when dealing with regression, it is prudent to begin with a smoother as a partial check on whether assuming a straight regression line appears to be reasonable. Simultaneously, the potential impact of outliers needs to be considered. In exploratory studies, it is suggested that results based on both Cleveland’s smoother and the running interval smoother be examined. (Quantile regression smoothers are another option that can be very useful; use the R function qsm.)

Here we begin by using the R function lplot (Cleveland’s smoother) to estimate the regression line when the goal is to estimate the mean of the dependent variable for some given value of an independent variable. Figure 9**A** shows the estimated regression line when using age to predict left corticospinal measures (CST.L). Figure 9**B** shows the estimate when a GORT fluency (GORT.FL) measure of reading ability (Wiederhold et al., 2001) is taken to be the dependent variable. The shaded areas indicate a 0.95 confidence region that contains the true regression line. In these two situations, assuming a straight regression line seems like a reasonable approximation of the true regression line.

**Figure 9:**
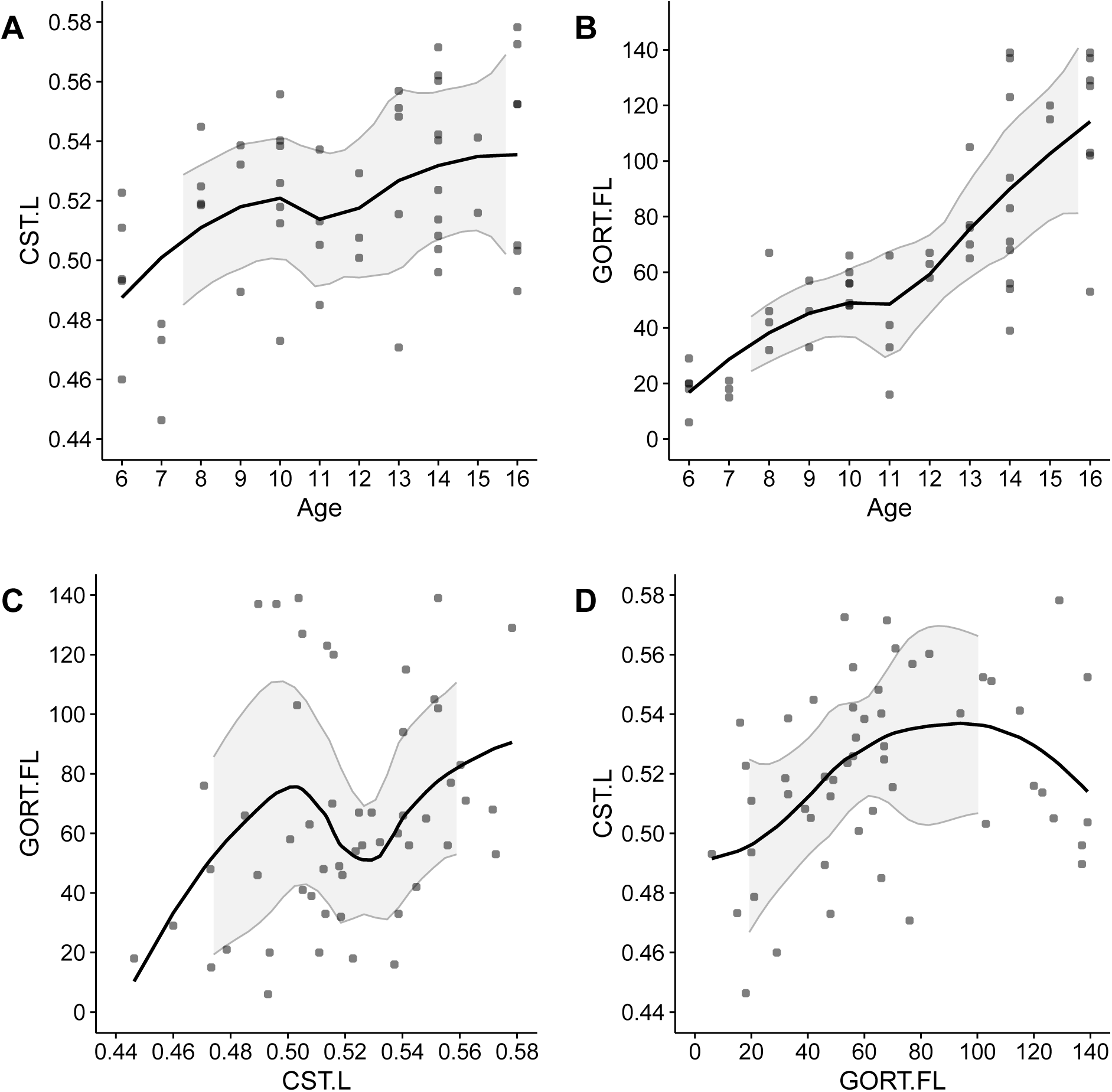
Non-parametric estimate of the regression line for predicting GORT.FL with CST.L

Figure 9**C** shows the estimated regression line for predicting GORT.FL with CST.L. Note the dip in the regression line. One possibility is that the dip reflects the true regression line, but another explanation is that it is due to outliers among the dependent variable. (The R function outmgv indicates that the upper GORT.FL values are outliers.) Switching to the running interval smoother (via the R function rplot), which uses a 20% trimmed mean to estimate the typical GORT.FL value, now the regression line appears to be quite straight. (For details, see the figshare file mentioned at the beginning of this section.)

The presence of curvature can depend on which variable is taken to be the independent variable. Figure 9**D** illustrates this point by using CST.L as the dependent variable and GORT.FL as the independent variable, in contrast to 9**C**. Note how the line increases and then curves down.

Using instead the running-interval smoother, where the goal is to estimate the 20% trimmed mean of the dependent variable, now there appears to be a distinct bend approximately at GORT.FL=70. For GORT.FL less than 70 the regression line appears to quite straight and the slope was found to be significant (p=0.007) based on the R function regci, which uses the robust Theil–Sen estimator by default. (It is designed to estimate the median of the dependent variable.) For GORT.FL greater than 70, again the regression line is reasonably straight, but now the slope is negative and does not differ significantly from zero (p=0.27). Moreover, testing the hypothesis that these two slopes are equal (via the R function reg2ci), p=0.027, which provides additional evidence that there is curvature. So a more robust smoother suggests that there is a positive association up to about GORT.FL=70, after which the association is much weaker and possibly nonexistent.

A common strategy for dealing with curvature is to include a quadratic term in the regression model. More generally, one might try to straighten a regression by replacing the independent variable *X* with *X^a^* for some suitable choice for *a*. However, for the situation at hand, the half slope ratio method (e.g., Wilcox, 2017a, 11.4) does not support this strategy. It currently seems that smoothers provide a more satisfactory approach to dealing with curvature.

Now consider the goal of determining whether age or CST.L is more important when GORT.FL is the dependent variable. A plot of the regression surface when both independent variables are used to predict GORT.FL suggests that the regression surface is well approximated by a plane. (Details are in the figshare document.) Testing the hypothesis that the two independent variables are equally important via the R function regIVcom indicates that age is more important. (This function uses the Theil–Sen estimator by default.) Moreover, the ratio of the strength of the individual associations is 53. Using least squares regression instead (via regIVcom but with the argument regfun=ols), again age is more important and now the ratio of the strength of the individual associations is 9.85.

Another measure of reading ability that was used is the Woodcock–Johnson (WJ) basic reading skills composite index (Woodcock, et al., 2001). Here we consider the extent the GORT (raw) rate score is more important than age when predicting the the WJ word attack (raw) score. Pearson’s correlation for the word attack score and age is 0.68 (p *<* 0.001). The correlation between the GORT rate score and the word attack score is 0.79 (p *<* 0.001). Comparing these correlations via the R function twohc4cor, no significant difference is found at the 0.05 level (p=0.11). But when both of these independent variables are entered into the model, and again the Theil–Sen regression estimator is used, a significant result is obtained (p< 0.001): the GORT rate score is estimated to be more important. In addition, the strength of the association between age and the word attack score is estimated to be close to zero. Using instead least squares regression, p = 0.02 and the correlation between age and WJ word attack score drops to 0.012. So both methods indicate that the GORT rate score is more important, with the result based on a robust regression estimator providing more compelling evidence that this is the case. This illustrates a point made earlier that the relative importance of the independent variables can depend on which independent variables are included in the model.

## 6 A Suggested Guide

While no single method is always best, the following guide is suggested when comparing groups or studying associations.

- Plot the data. Error bars are popular, but they are limited regarding the information they convey, regardless of whether they are based on the standard deviation or an estimate of the standard error. Better are scatterplots, boxplots or violin plots. For small sample sizes, scatterplots should be the default. If the sample sizes are not too small, plots of the distributions can be very useful, but there is no agreed upon guideline regarding just how large the sample size should be. For the moment, we suggest checking both boxplots and plots of the distributions when the sample size is *n* ≥ 30 with the goal of getting different perspectives on the nature of the distributions. It is suggested that kernel density estimators, rather than histograms, be used for reasons illustrated in Wilcox (2017b). (The R functions akerd and g5plot use a kernel density estimator.) For discrete data, where the number of possible values for the outcome variable is relatively small, also consider a plot of the relative frequencies. The R function splot is one possibility. When comparing two groups consider the R function splotg2. For more details regarding plots, see Weissgerber et al. (2015) and Rousselet et al. (2017).
- For very small sample sizes, say less than or equal to ten, consider the methods in sections 4.2 and 4.3.
- Use a method that allows heteroscedasticity. If the homoscedasticity assumption is true, in general little is lost when using a heteroscedastic method. But as the degree of heteroscedasticity increases, at some point methods that allow heteroscedasticity can make a practical difference in terms of both Type I errors and power. Put more broadly, avoid methods that use an incorrect estimate of the standard error when groups differ or when dealing with regression and there is an association. These methods include t-tests and ANOVA F tests on means, the WMW test, as well as conventional methods for making inferences about measures of association and the parameters of the usual linear regression model.
- Be aware of the limitations of methods based on means: they have a relatively high risk of poor control over the Type I error probability, as well as poor power. Another possible concern is that when dealing with skewed distributions, the mean might be an unsatisfactory summary of the typical response. Results based on means are not necessarily inaccurate, but relative to other methods that might be used, there are serious practical concerns that are difficult to address. Importantly, when using means, even a significant result can yield a relatively inaccurate and unrevealing sense of how distributions differ, and a non-significant result cannot be used to conclude that distributions do not differ.
- As a useful alternative to comparisons based on means, consider using a shift function or some other method for comparing multiple quantiles. Sometimes these methods can provide a deeper understanding of where and how groups differ that has practical value as illustrated in Figure 7. There is even the possibility that they yield significant results when methods based on means, trimmed means and medians do not. For discrete data, where the variables have a limited number of possible values, consider the R function binband. (See Wilcox, 2017c, section 12.1.17; or Wilcox, 2017a section 5.8.5.)
- When checking for outliers, use a method that avoids masking. This eliminates any method based on the mean and variance. Wilcox (2017a, section 6.4) summarizes methods designed for multivariate data. (The R functions outpro and outmgv use methods that perform relatively well.)
- Be aware that the choice of method can make a substantial difference. For example, highly non-significant results can become significant when switching from a method based on the mean to a 20% trimmed mean or median. The reverse can happen where methods based on means are significant but robust methods are not. In this latter situation, the reason might be that confidence intervals based on means are highly inaccurate. Resolving whether this is the case is difficult at best based on current technology. Consequently, it is prudent to consider the robust methods outlined in this paper.
- When dealing with regression or measures of association, use modern methods for checking on the impact of outliers. When using regression estimators, dealing with outliers among the independent variables is straightforward via the R functions mentioned here: set the argument xout=TRUE. As for outliers among the dependent variable, use some robust regression estimator. The Theil–Sen estimator is relatively good, but arguments can be made for using certain extensions and alternative techniques. When there are one or two independent variables, and the sample size is not too small, check the plots returned by smoothers. This can be done with the R functions rplot and lplot. Other possibilities and their relative merits are summarized in Wilcox (2017a).

## 7 Concluding Remarks

It is not being suggested that methods based on means should be completely abandoned or have no practical value. Instead, complete reliance on conventional methods can result in a superficial, misleading and relatively uninformative understanding of how groups compare. In addition, they might provide poor control over the Type I error probability and power. Similar concerns plague least squares regression and Pearson’s correlation.

When a method fails to reject, this leaves open the issue of whether a significant result was missed due to the method used. From this perspective, multiple tests can be informative. However, there are two competing goals that need to be considered. The first is that when testing multiple hypotheses, this can increase the probability of one or more Type I errors. From basic principles, if, for example, five tests are performed at the 0.05 level, and all five hypotheses are true, the expected number of Type I errors is 0.25. But the more common focus is on the probability of one or more Type I errors rather than the expected number of Type I errors. The probability of one or more Type I errors will be greater than 0.05, but by how much is difficult to determine exactly due to the dependence among the tests that are performed. Improvements on the Bonferroni method deal with this issue (e.g., Hochberg, 1988; Hommel, 1988) and are readily implemented via the R function p.adjust. But the more tests that are performed, such adjustments come at the cost of lower power. Simultaneously, ignoring multiple perspectives runs the risk of not achieving a deep understanding of how groups compare. Also, as noted in the previous section, if methods based on means are used, it is prudent to check the extent robust methods give similar results.

An important issue not discussed here is robust measures of effect size. When both conventional and robust methods reject, the method used to characterize how groups differ can be crucial. Cohen’s *d*, for example, is not robust simply because it is based on the means and variances. Robust measures of effect size are summarized in Wilcox (2017a, c). Currently, efforts are being made to extend and generalize these measures.

Another important issue is the implication of modern insights in terms of the massive number of published papers using conventional methods. These insights do not necessarily imply that these results are incorrect. There are conditions where classic methods perform reasonably well. But there is a clear possibility that misleading results were obtained in some situations. One of the main concerns is whether important differences or associations have been missed. Some of the illustrations in Wilcox (2017a, b, c), for example, reanalyzed data from studies dealing with regression where the simple act of removing outliers among the independent variable resulted in a highly non-significant result becoming significant at the 0.05 level. As illustrated here, non-significant results can become significant when using a more modern method for comparing measures of central tendency. There is also the possibility that a few outliers can result in a large Pearson correlation when in fact there is little or no association among the bulk of the data (Rousselet & Pernet 2012). Wilcox (2017c) mentions one unpublished study where this occurred. So the issue is not whether modern robust methods can make a practical difference. Rather, the issue is how often this occurs.

Finally, there is much more to modern methods beyond the techniques and issues described here (Wilcox, 2017a). Included are additional methods for studying associations as well as substantially better techniques for dealing with ANCOVA. As mentioned in the introduction, there are introductory textbooks that include the basics of modern advances and insights. But the difficult task of modernizing basic training for the typical neuroscientist remains.

